# Colony size frequency distribution across gradients of reef health in disturbed coral reefs in Northeast Peninsular Malaysia

**DOI:** 10.1101/2022.05.21.492910

**Authors:** Gilles Gael Raphael Bernard, Alexandra Lucie Kellam, Sebastian Szereday

## Abstract

Coral reefs globally are experiencing chronic stress leading to the deterioration of health and functionality. Analysis of size frequency distribution (SFD) of hard corals enables post hoc assessments of major demographic events (e.g., recruitment and mortality) that follow ecological disturbances. Here, we present an evaluation of current reef health, SFD and recruitment of 37 morpho-taxa in Northeast Peninsular Malaysia. Results highlight stress viable demographic structure of massive taxa (e.g., massive *Porites*) and significant differences of SFD across gradients of reef health, whereby degraded sites were predominantly characterized by negatively skewed (e.g., large colonies) and leptokurtic (e.g., high population turnover) distribution of dominant hard coral taxa. Ultimately, results suggest that locally coral reef degradation can exceed tipping points, after which annual monsoon conditions and degraded reef substrates interact to reinforce and manifest negative feedback loops, thereby impeding demographic recovery, and altering coral SFD and population assemblage.

## 1. Introduction

Coral reefs have entered a crucial decade in which climate change associated extremes (e.g., global ocean warming and cyclones) are posing an existential threat to coral reef survival worldwide (Hughes et al. 2017a, 2017b). Multiple, synchronous and synergistic disturbances of natural (e.g., seasonal storms) and human origin (e.g., overfishing, nitrification, coastal development) are having a direct impact on coral reefs worldwide and their population viability (Hughes and Connell 1999; Bellwood et al., 2004; Harborne et al., 2016). Population dynamics, function and resilience of ecosystems are co-determined by the reproductive viability of a population, driven by the rates of births and deaths, and the amount of sexually mature individuals (Weinstein and Pillai, 2015). Inherent demographic variations across time and space are the results of events altering these processes, whereby present-day demographic structures can hold clues over time and extent of past system-wide perturbations (Swedlund, 1978). In clonal organisms, such as hard corals, responses and adaptations to environmental disturbances are reflected in the size frequency distribution (SFD) of individual populations (Bak and Meesters, 1998), and these often vary greatly among taxonomic groups (Meesters et al., 2001). Moreover, size-dependent mortality following large-scale disturbances such as coral bleaching and tropical storms (Loya et al., 2001; Baird and Marshall, 2002) likely dictates shifts in population assemblage and size structure (Massel and Done, 1993; McClanahan et al., 2001; Anderson and Pratchett, 2014). Modifications in the distribution and frequency of particular size groups can assert major changes in hard coral community assemblages with cascading impacts on the structural habitat of reef species, and therefore, on the functional and economic value of coral reefs (Graham, 2014; Darling et al., 2017). Thus, disturbance intensity and frequency, as well as taxon-identity and population size structure co-determine the resilience and trajectory of coral reefs (Bak and Meesters, 1998, 1999; McClanahan et al., 2008; Darling et al., 2013; Dietzel et al. 2020).

The study of age-size relationships of sedentary organisms such as trees (dendrology) has been well-established to identify fluctuations and disturbances in the biophysical environment (Fonti et al., 2009; Loader et al., 2010). Similar methods (e.g., sclerochronology) have been applied to coral reef research to reconstruct historic environmental variability (Barnes, 1970; Hudson et al., 1976; Lough and Cantin, 2014). However, identifying age-size relationships in scleractinian corals remains challenging due to fragmentation and partial mortality of individual colonies, as well as fusion of genetically different colonies (Hughes and Jackson, 1980; Babcock, 1991). The scleractinian demographic structure and SFD is highly variable across fine spatial scales such as individual reefs and across multiple depths (Adjeroud et al., 2007, 2015; Kramer et al., 2020), but is an appropriate method to study scleractinian population dynamics and status in field-assessments (Hughes and Connell, 1987). Indeed, vital functional traits that underpin the functioning and resilience of coral reefs, such as fecundity, growth, partial and total mortality (Darling et al., 2012), are influenced by colony size rather than age (Hughes and Jackson, 1985; Hall and Hughes 1996). Therefore, colony SFD is suitable to compare inter- and intra-specific variation in community size structure as to provide insights into past ecosystem wide disturbance events (Bak and Meesters, 1998), and to elaborate on demographic strategies of scleractinian taxa that underline changes in coral cover and reef health (Miller et al., 2016). Ultimately, SFD assessments are viable tools to estimate the impacts of multiple and synchronous stressors on scleractinian assemblages and populations (Fong and Glynn, 1998; Smith et al., 2005), particularly when historical and qualitative data sets are absent and parochial. Considering the multitude of present stressors on coral reefs (Hughes and Connell, 1999; Bellwood et al., 2004; Hughes et al., 2017b), as well as the lack of demographic assessments in field studies, SFD and demographic structure assessments are urgently required to understand ecological changes in hard coral communities under continuous environmental stress (Edmunds and Riegl, 2020).

In 2003, the average coral cover in the Indo-Pacific was reduced to approximately 22 %, after decreasing annually by 1-2 % between the 1980s and 2003 (Bruno and Selig, 2007). However, the percent coral cover metric is not conclusive and cannot detect significant changes in the capability of coral reefs to recover and maintain ecosystem functioning (Hughes et al., 2010; Edmunds & Riegl, 2020), as shifts in cover are underlined by changes in demographic and size structure (Miller et al., 2016; Dietzel et al., 2020). Despite high regional diversity of hard coral species (Huang et al., 2015), and high economic value of coral reefs (Kamarruddin et al., 2013; Sukarno et al., 2015), data on Malaysian hard coral reefs is limited to benthic cover assessments with a focus on hard coral reef coverage and taxonomic richness (Harborne et al., 2000). Qualitative and quantitative information on hard coral communities, particularly coral recruitment, general demographic studies and studies detailing community size structure, are distinctively lacking or are unavailable in Malaysia (Praveena et al., 2012). As coral reefs in Peninsular Malaysia are in a state of decline (Toda et al., 2007; Reef Check Malaysia, 2019), studies are required to determine stochastic shocks and pulses that underpin population dynamics and recovery, such as overall demographic structure (e.g., ratio of coral recruitment and sexually mature individuals). In view of limited historical data and in consideration of current disturbance regimes, determining the size-frequency distributions can serve as a post hoc assessment of demographic events (mortality, recruitment, population turnover), to provide insights into demographic shifts and to elaborate future trajectories.

Two theories have been suggested to explain shifts and non-equilibria states of hard corals demographics. Firstly, a transition towards relatively more abundance of large colonies has been demonstrated in the Caribbean (Bak and Meesters, 1999; Meesters et al., 2001; Miller et al., 2016) and the Great Barrier Reef (Dietzel et al., 2020), particularly post multiple large-scale disturbance events. In contrast, a shift towards relatively more small colonies has been documented in French Polynesia (Adjeroud et al., 2015), in the western Indian Ocean (McClanahan et al., 2008), and in the in the Red Sea (Riegl et al., 2012), whereby constant impulses of coral recruitment were suggested to further drive shifts in SFD in the Red Sea and partially in French Polynesia. Moreover, persistent ecosystem-wide disturbance shocks and chronic stress exposure systematically narrow colony size ranges and homogenize community size structures (Cannon et al., 2021). Consequently, this study investigated 1.) SFD and demographic structure in Northeast Peninsular Malaysia across a continuum of reef health (e.g., cover, diversity, density, etc.), which results from biophysical reef site conditions (e.g., leeward vs windward), and negative impacts of anthropogenic origin (sewage, coastal development, overfishing and rising SST). Secondly, we investigated 2.) present patterns of coral recruitment, to determine whether recent stochastic impulses of coral recruitment possibly resulted in the present demographic structure (Riegl et al., 2012). Ultimately, these findings highlight 3.) hard coral assemblages and taxa that are potentially more tolerant of multiple stressors and are more likely to persist under current and future scenarios of persistent (human) impacts. Hereby, post hoc evidence for the second hypothesis (shift to smaller colonies) should highlight recently established populations (e.g., after disturbance) of any given taxa with a homogenous size structure, composed of smaller colonies and a peaked distribution (*sensu* McClanahan et al., 2008; Riegl et al., 2012), whereas potentially stress tolerant taxa should present with a more evenly distributed spectrum of size classes (*sensu* Adjeroud et al., 2015). Additionally, reefs at advanced stages of degradation should be characterized by highly centralized distribution of predominantly weedy taxa and a preponderance of large colonies (e.g., disturbance ‘survivors’; *sensu* Meesters et al., 2001; Dietzel et al., 2020). Lastly, intra-specific differences in SFD should be well defined across gradients of reef health, (*sensu* Bauman et al., 2013), to accurately highlight whether taxon identity or reef conditions underline the hard coral SFD. This study represents the first analysis of the demographic structure and SFD of scleractinian taxa in Peninsular Malaysia to date and provides fundamental insights to guide future management of hard coral reefs in Peninsular Malaysia.

## 2. Methods

### 2.1 Study location

Fieldwork and data collection was carried out between September and October 2019 around Pulau Lang Tengah (Pulau=Island) in Northeast Peninsular Malaysia (5°47’49.7”N, 102°53’45.0”E) (Figure 1). Pulau Lang Tengah is included in the Pulau Perhentian and Pulau Redang marine park, and is legally protected from resource extraction and fishing (Praveena et al., 2012). However, monitoring of fishing activities and enforcement of policies is limited (Kimura et al., 2022). Despite legal protection from fishing pressure, over- and illegal fishing is a significant problem in Malaysia (Stodbuzki et al., 2006, Asia-Pacific Economic Cooperation, 2008; Ghazali et al., 2019). Low numbers of herbivorous reef fish have been documented around Pulau Lang Tengah, whereby total herbivorous fish abundance was lowest at leeward sites (Chew, 2018, unpublished data), suggesting elevated pressure on leeward coral reef resilience and colony size in view of reduced herbivorous grazing (McClanahan et al., 2001; Green and Bellwood, 2009). Thus, three sites along a set of human stressors and along a wind gradient were selected: Pasir Besar (PB), Batu Kucing (BK) and Batu Bulan (BB), whereby BB and BK are windward facing fringing reefs, and PB is a leeward lagoon (Figure 1A-B). Wind frequency and direction were determined by using web-based monitoring products (Global Wind Atlas 3.0). In addition, all sites are subjected to the annual Northeast Monsoon (November to February), which intensifies hydrodynamic conditions and increases wave intensity; factors that are known to consequentially influence scleractinian colony SFD (Madin and Connolly, 2006; Madin et al., 2012, 2014). Moreover, the leeward shore has been extensively developed (Figure 1A), supposedly resulting in physical degradation of adjacent coral reefs (Praveena et al., 2012), as well as untreated sewage discharge from nearby resorts, further pressuring leeward reef assemblages (Wooldridge et al., 2012; Reef Check Malaysia, 2019). Physical reef degradation may further result in larger quantities of mobile substrate (e.g., coral rubble) which impede natural reef recovery due to sedimentation and physical interference (Fox et al., 2019; Wolfe et al., 2021). Ultimately, there is a clear delineation of local site pressures with markedly higher negative anthropogenic impacts on leeward coral reefs, steering from the close proximity of leeward coral reefs to the source of the disturbance (e.g., beach resorts) and subsequent secondary impacts (Fisher et al., 2008). Lastly, regional ecosystem wide disturbance events that likely impacted scleractinian SFD at all sites, such as coral bleaching (McClanahan et al., 2001, 2008), have been recorded across the entire east coast of the Malayan Peninsula in 1998 (Kushairi, 1998), in 2010 (Tan and Heron, 2011; Guest et al., 2012) and in May 2019 at all sites around Pulau Lang Tengah before data collection (Szereday and Affendi, 2022). Moreover, tropical cyclone Pabuk significantly reduced live hard coral cover in January 2019 in Northeast Peninsular Malaysia (Reef Check Malaysia, 2019; Safuan et al., 2020), suggesting probable impacts on hard coral SFD regionally. Subsequently, any observed differences in SFD and general coral reef health may partially be the result of synergistic impacts from coral bleaching (e.g., 1998 and 2010), hydrodynamic disturbances (e.g., cyclone Pabuk 2019, annual monsoon), sedimentation (amplified by physical degradation after coastal development), overfishing, eutrophication and fundamental differences in hard coral assemblages and colony spatial-competition. Therefore, our deliberate study design attempted to highlight differences in hard coral SFD across a continuum of reef health and under various sets of local and regional stressor agents. This determined focus on site differences (e.g., comparing degraded to healthier reef sites) is essential as historical data on coral reef health and population size structure are unavailable for Northeast Peninsular Malaysia, and baseline comparisons to studies prior to the occurrence of mass disturbances is not possible. Hereby, it is important to note that healthier reef sites do not necessarily reflect pristine conditions of former reefs, and this study represents a post hoc analysis of possible demographic events that steered from human disturbances.

**Figure 1.**
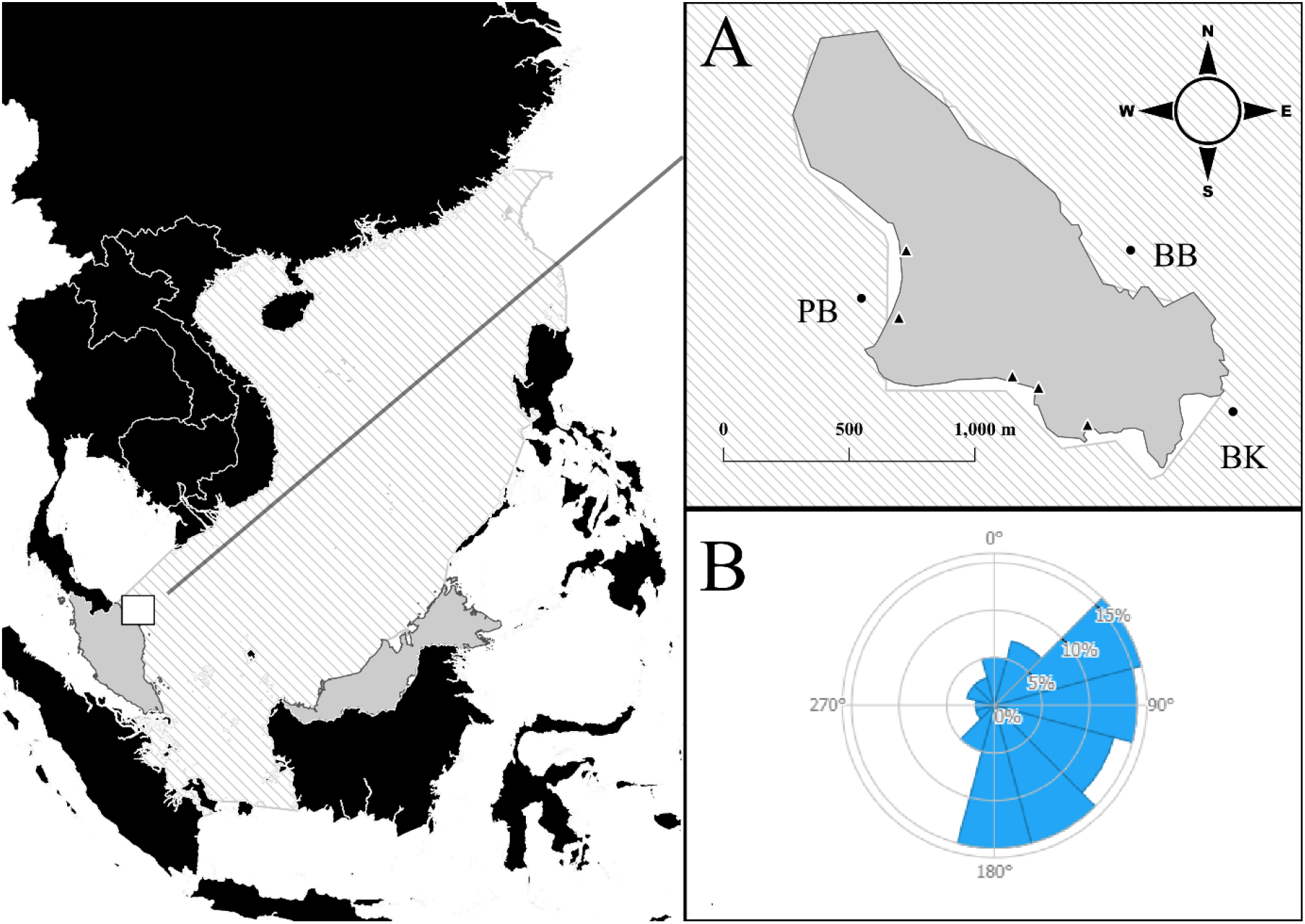
Location of Pulau Lang Tengah (A) in Northeast Peninsular Malaysia and within the South China Sea basin (dashed line area). Survey sites are shown by circles and abbreviations (PB – Pasir Besar; BK – Batu Kucing; BB-Batu Bulan). A wind frequency rose (B) illustrates wind direction and frequency at island scale. Locations marked with triangles along the shoreline demarcate coastal development sites (e.g., beach resorts).

### 2.1. Benthic surveys and reef health indicators

At each site, benthic surveys were conducted along three 50 meters transects (measure tapes) laid parallel to the shore (Figure 1) and across multiple depths: shallow (4-8 m), intermediate (8-12 m) and deep (15-20 m). Transect lines were anchored with weights to ensure maximum tape stretch and to follow the reef contour. Surveys were performed by three divers in equal proportion to reduce observer bias. Each transect consisted of one 50 meters line intercept transect (LIT) and one 30 x 1 m belt transect. To reduce spatial sampling biases, belt transects were broken up into six 5 x 1 m subsets at randomized points along the 50 m transect, and were split equally across both sides of the tape (total of 15 x 1 m per side). Belt transect data was used to determine scleractinian colony size frequency distribution and various reef health indicators (e.g., density and diversity), whereas LITs were used to measure benthic substrate composition in order to further approximate reef health (Teichberg et al., 2018). Hereby, non-living substrate included sand, rock, recently killed corals and coral rubble, and following live-benthic substrates were recorded: hard corals, soft corals, sponges, sea anemones, algae, giant clams, zoanthids and corallimoprhs, as well as smaller benthic organisms (e.g., organ pipe corals, fern corals, etc.). All living benthic substrates recorded during LITs were measured from the colonies edge-to-edge and to the nearest centimetre, occasionally requiring the use of a reference stick to perpendicularly project the colony edges onto the measure tape. This resulted in complete measurements of colony length along the substrate rather than shortened intercept measurements, whereas non-living substrates were recorded to the nearest centimetre by measuring the intercept length. Subsequently, the sum of all recorded lengths along the transect tape exceeded the actual 50 meters transect length. Therefore, following formula was applied to calculate percent benthic coverage of a benthic substrate group (e.g., hard coral cover), where *TL* is the standard 50 meters transect length:

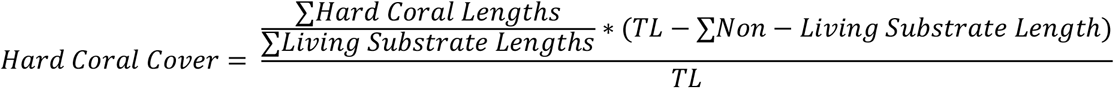

Note, hard coral cover includes the percent cover of *Heliopora spp*. colonies, due to their reef building function on Indo-Pacific reefs. (Colgan, 1984). Individual hard coral colonies were defined as autonomous, free-standing coral colonies with live tissue. Autonomous colonies with fragmented live tissue resulting from partial colony mortality were considered a single colony (Bak and Meesters, 1998), and the amount of partial mortality was visually estimated in steps of 5% relative to the total colony surface area. Lastly, LIT measurements of hard coral colony length were used to measure minimum and maximum colony length in addition to size class that were recorded during belt transects.

Further indicators of reef health, such as taxonomic richness, evenness and colony density of various size groups as well as colony size (Fisher et al., 2008), were calculated for each site based on belt transect data to highlight coral reef health across sites. Belt transects increased sampling size, accuracy, and reduced sampling errors of LITs, which are not suitable to detect very small colonies (e.g., < 3 cm) (Obura and Grimsditch, 2009). Along belt transects, the size of each hard coral colony encountered (including octocoral *Heliopora spp*.), was recorded by measuring their maximum horizontal and linear extension with a graduated reference pipe. Colony size classes were based on Obura and Grimsdtich (2009): 0-2.5 cm, 2.5-5.0 cm, 5-10 cm, 10-20 cm, 20-40 cm, 40-80 cm, 80-160 cm, 160-320 cm and colonies > 320 cm. Here, colonies in the two smallest size classes (< 5cm) were considered recruits. Furthermore, to reduce Type 1 and Type 2 sampling errors (Zvuloni et al., 2008), only colonies whose centre was within the belt were recorded (Nugues and Roberts, 2003), in addition to colonies with at least 40% of colony surface area inside the belt.

Taxonomic identification was performed using the Coral Finder 3.0 (Kelley, 2016), and the unique morphological appearances of individual colonies were recorded. Differences in SFD and demographic structure can be detected on taxon level, and greater taxonomic resolution may lead to more pronounced differences. However, correct identification of species during in-water surveys based on morphological characteristics remains challenging. Therefore, to obtain optimal resolution while minimizing identification biases, survey analysis of SFD and demographic structure was conducted on distinctive morphological and taxonomic levels (e.g., *massive Porites*, *encrusting Porites*), henceforth referred to as morpho-taxon. This resolution is suitable to describe demographic structure and SFD driven by taxon identity and functional traits (growth, fecundity, etc.), which are phylogenetically conserved on genus and morphological level (Darling et al., 2012; Alvarez-Noriega et al., 2016). In addition, *Porites* species with an encrusting plate and up-growths (exclusively compromising *Porites rus* and similar species), were grouped together due to their distinctive morphology and are hereafter referred to as *Porites spp. (rus)*. Comparison of SFD on morphological level was based on seven groupings: branching forms (e.g. arborescent, digitate, corymbose, hispidose), massive forms (massive and sub-massive), encrusting (flat crust), encrusting with up-growths (e.g., crusts with short vertical branches and with columns), laminar forms (e.g. vase, plate, foliose), columnar colonies and solitary (free-living) colonies.

### 2.2. Size frequency distribution (SFD) and statistical analysis

Data and statistical analyses were performed in R software (R Core Team, 2020). Prior to analysis, non-normality of the SFDs was formally tested with Shapiro-Wilk tests (SW, package ‘RVAideMemoire’) (Hervé, 2021). To reduce non-normality, colony sizes were log_10_ transformed by using the middle value of each size class (the middle value used for the size class > 320 cm was 480 cm). The log transformed SFD distribution of morphological groups and morpho-taxa was compared among sites and depths. Hereby, we used Anderson-Darling (AD) k-samples tests (package ‘kSamples’) (Scholz and Zhu, 2019), to compare intra-morphological and intra-taxonomic SFD differences across sites. Anderson-Darling (AD) k-samples tests were conducted with the assumption that all samples came from the same distribution, and p-values were adjusted with the Bonferroni correction method (Abdi, 2017). Moreover, the AD test was chosen as it is a non-parametric test, it reliably detects small variations, it is applicable to discrete distributions, and requires less data to reach sufficient statistical power in order to detect differences between varying sample sizes. (Engmann and Cousineau, 2011). As size classes are non-continuous, the second version was used with 10,000 simulations based on random splits of the pooled samples to estimate the exact conditional p values. Hereby, for each computed analysis, a minimum sample size of seven colonies was applied. Significant differences in scleractinian colony size were explored using non-parametric Kruskal-Wallis tests (KW, package ‘stats’, R Core Team, 2021) to explore differences within groups (e.g., site specific morphological groups), and Mann-Whitney-Wilcoxon tests (MW, package ‘stats’) to explore differences between two sampling populations (e.g., morpho-taxonomic comparisons). The distribution curves of SFD were then analysed with descriptive statistical measures to investigate skewness (g_1_) and kurtosis (g_2_) (package ‘moments’) (Komsta and Novomestky, 2015). Here, positive skewness identifies a dominance of small colonies and a negatively skewed distribution results from predominance of large colonies. Kurtosis identifies the peakedness of a distribution near its central mode and highlights whether the distribution tails contain extreme values. A peaked distribution is leptokurtic (g_2_ > 0; dominance of a few consecutive size classes), and a distribution flatter than a normal distribution is platykurtic (g_2_ < 0; higher variation in size class abundance). Skewness and kurtosis were divided by their respective standard errors to determine whether the values were significantly different from the normal distribution (Cramer, 1997; Wright and Herrington, 2011):

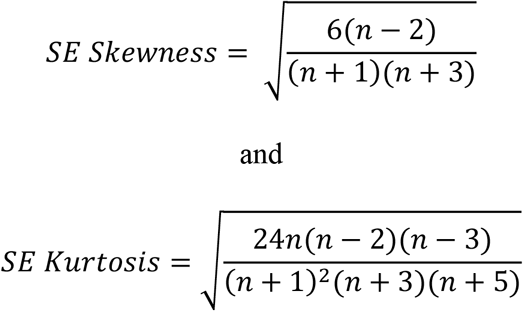

Here, a distribution was significantly different from the normal distribution if the Z-value of the skewness and kurtosis statistic was > ±2. Moreover, variations in colony size were expressed by the coefficient of variation (CV), low values suggesting low variation in colony size, and we further quantified the 10^th^ and 90^th^ percentile of each log-transformed colony size. Finally, differences in morpho-taxonomic assemblages across sites and depths were visualized using a non-metrical multidimensional scaling (NMDS) visualization in a Bray-Curtis dissimilarity matrix, modelled with the package ‘vegan’ (Oksanen et al., 2020) in R software.

## 3. Results

### 3.1. Benthic surveys and reef health

Benthic surveys revealed substantial difference in coral assemblages across sites and depth (Figure 2). All reef health indicators including hard coral density, diversity and percent hard coral cover were consistently lower at leeward PB (Table 1), and increased along the reef health and wind gradient from PB (leeward, least wind exposed) to BK and BB (most wind exposed). Accordingly, hard coral cover was lowest at PB (8.78%), followed by BK (14.94%) and was highest at BB (43.37%). Consistent with the hard coral cover gradient from PB to BB, benthic cover at leeward PB (48.11 % sand and 39.61 % coral rubble) and windward BK (41.87 % sand and 32.03 % coral rubble), was markedly more dominated by non-living substrate compared to windward BB (14.37 % sand and 11.82 % coral rubble). At island scale, benthic cover was dominated by sand (34.78 %) and rubble (27.82 %), and island wide hard coral cover was 22.26 %. Regardless of sites, hard coral cover declined with increasing depth (Supplementary S1). Island wide hard coral density corresponded to 15.06 (colonies / m^2^), following the same gradient as hard coral cover, with the lowest density at leeward PB (9.21 colonies / m^2^), to windward BK (10.26 colonies / m^2^), and BB (highest density, 25.71 colonies / m^2^). The Shannon Index of morpho-taxa increased from leeward PB to BK and BB (2.45, 2.70 and 3.15, respectively).

**Figure 2.**
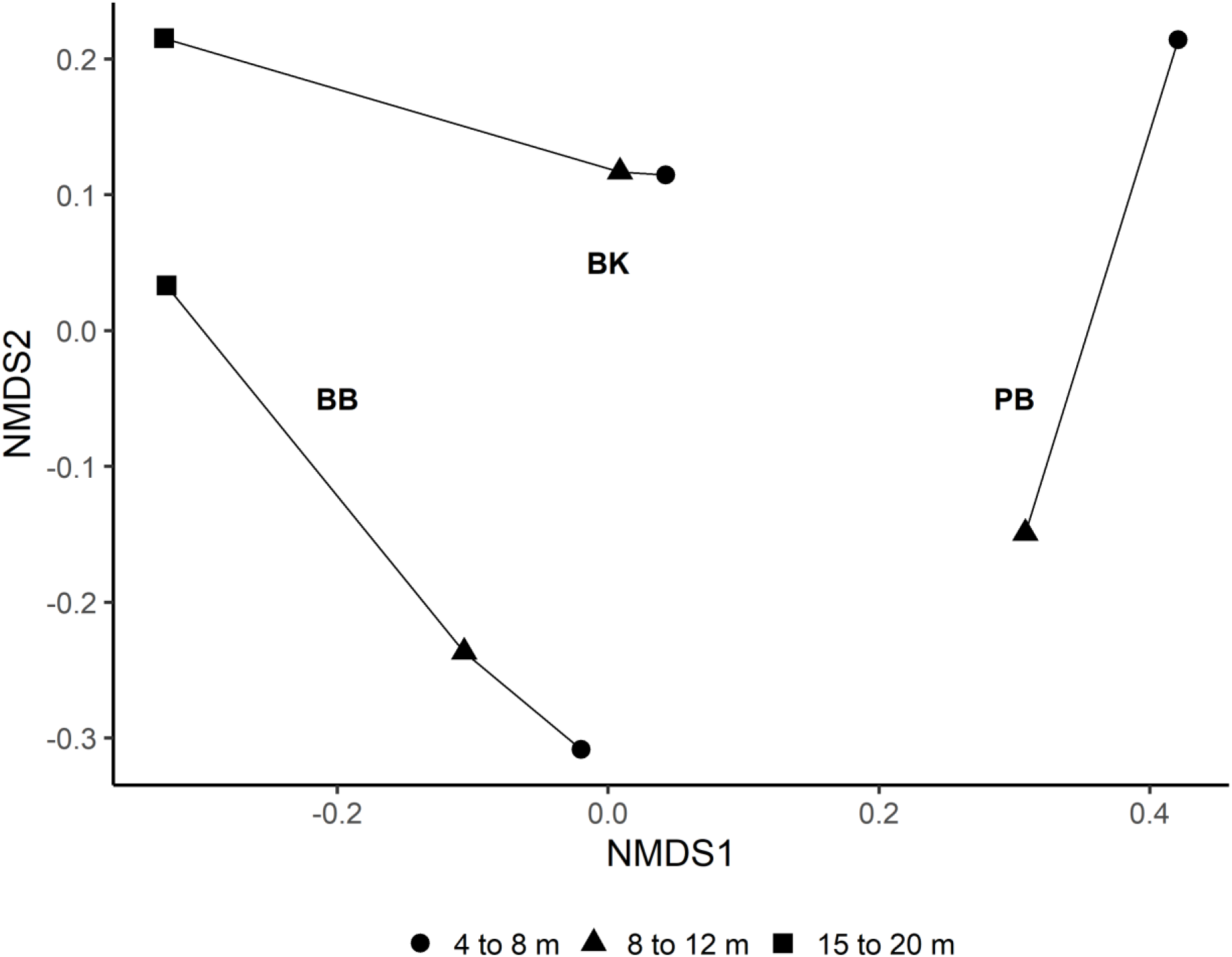
Hard coral community assemblage similarity is presented as a function of depth and site in a non-metric multidimensional scaling (NMDS) ordination plotted on Bray-Curtis dissimilarity matrices (2D). Each line represents a survey site (BB-Batu Bulan; BK-Batu Kucing, PB-Pasir Besar) and markers present the morpho-taxnomic assemblage at the respective site and depth, where close proximity of markers highlights similarities in hard coral community assemblages. Note, no hard coral colonies were recorded at 15 to 20 meters of depth at Pasir Besar (PB).

**Table 1.** Summary and comparison of benthic substrates, coral reef health indicators and demographic variables of hard corals across survey sites around Pulau Lang Tengah, Northeast Peninsular Malaysia. Batu Bulan and Batu Kucing are windward facing sites, and Pasir Besar is leeward facing. Highlighted in grey are demographic variables that are significantly different from the mean distribution.

|  | Lang Tengah | Batu Bulan | Batu Kucing | Pasir Besar |
| --- | --- | --- | --- | --- |
| Number of genera | 40 | 33 | 30 | 24 |
| Number of morpho-taxa | 72 | 58 | 46 | 38 |
| Number of colonies (n) | 4066 | 2314 | 923 | 829 |
| Shannon Index (H) | 3.15 | 3.17 | 2.70 | 2.45 |
| Shannon Index (H) recruits (<5 cm) | 3.06 | 3.10 | 2.63 | 2.60 |
| Recruit colonies / m<sup>2</sup> (<5 cm) | 5.56 | 9.40 | 4.42 | 2.87 |
| Sub-adults colonies / m<sup>2</sup> (≥5-20 cm) | 7.09 | 11.61 | 4.38 | 5.27 |
| Adult colonies / m<sup>2</sup> (>20-80 cm) | 2.23 | 4.37 | 1.28 | 1.03 |
| Large colonies / m<sup>2</sup> (>80 cm) | 0.19 | 0.33 | 0.18 | 0.04 |
| Mean colony size<sub>log</sub> cm (SD) | 0.87 (0.48) | 0.88 (0.48) | 0.80 (0.50) | 0.89 (0.42) |
| Colony size coefficient of variation % (CV) | 54.79 | 54.48 | 62.59 | 46.68 |
| Skewness (g<sub>1</sub>) (Z-value) | 0.03 (0.78) | 0.07 (1.38) | **0.19 (2.36)** | **-0.35 (-4.13)** |
| Kurtosis (g<sub>2</sub>) (Z-value) | **-0.39 (-5.09)** | **-0.40 (-3.94)** | **-0.47 (-2.94)** | -0.25 (-1.48) |
| % hard coral cover | 22.26 | 43.37 | 14.94 | 8.78 |
| % other living benthos | 4.94 | 12.61 | 0.90 | 0.99 |
| % coral rubble | 27.82 | 11.82 | 32.03 | 39.61 |
| % sand | 34.78 | 14.37 | 41.87 | 48.11 |
| % dead hard corals | 2.45 | 4.34 | 1.39 | 1.63 |

### 3.2. Demographic structure and recruitment

Sub-adult colonies as well as coral recruits dominated the screlactinian demographic structure around Pulau Lang Tengah, whereby sub-adults accounted for 47.08% (n=1,914) and recruits for 36.94% (n=1,502) of surveyed colonies (n=4,066). This general dominance of small sized colonies was further highlighted by the low abundance (0.19 colonies / m^2^) of large colonies (>80 cm) and was largely consistent across sites (Table 1). However, more intact sites hosted a higher amount of large and small colonies, and the mean colony size of morpho-taxa was different across taxa and sites, whereby the coefficient of variations of colony size varied from 62.59% at BK, 54.48% at BB and 46.68% at PB (Table 1). Secondly, there were notable variations in the abundance of coral recruits across sites (Supplementary S2), whereby the relative recruitment density was substantially higher at windward BB (9.40 recruits / m^2^) than at BK (4.42 recruits / m^2^) and PB (2.87 recruits / m^2^) (Table 1). Furthermore, coral recruitment was disproportionately dominated by few taxa (Figure 3). On morpho-taxonomic level, recruits of massive *Porites* (18.0% of total recruitment, 3.01 recruit / m^2^) and *Porites spp. (rus)* (9.5%, 1.59 recruits / m^2^) were most abundant, followed by massive *Favia* (8.9%, 1.48 recruits / m^2^), solitary corals *Fungia* (7.3%, 1.22 recruits / m^2^), and encrusting *Leptastrea* (6.0%, 1.00 recruits / m^2^) (Figure 3).

**Figure 3.**
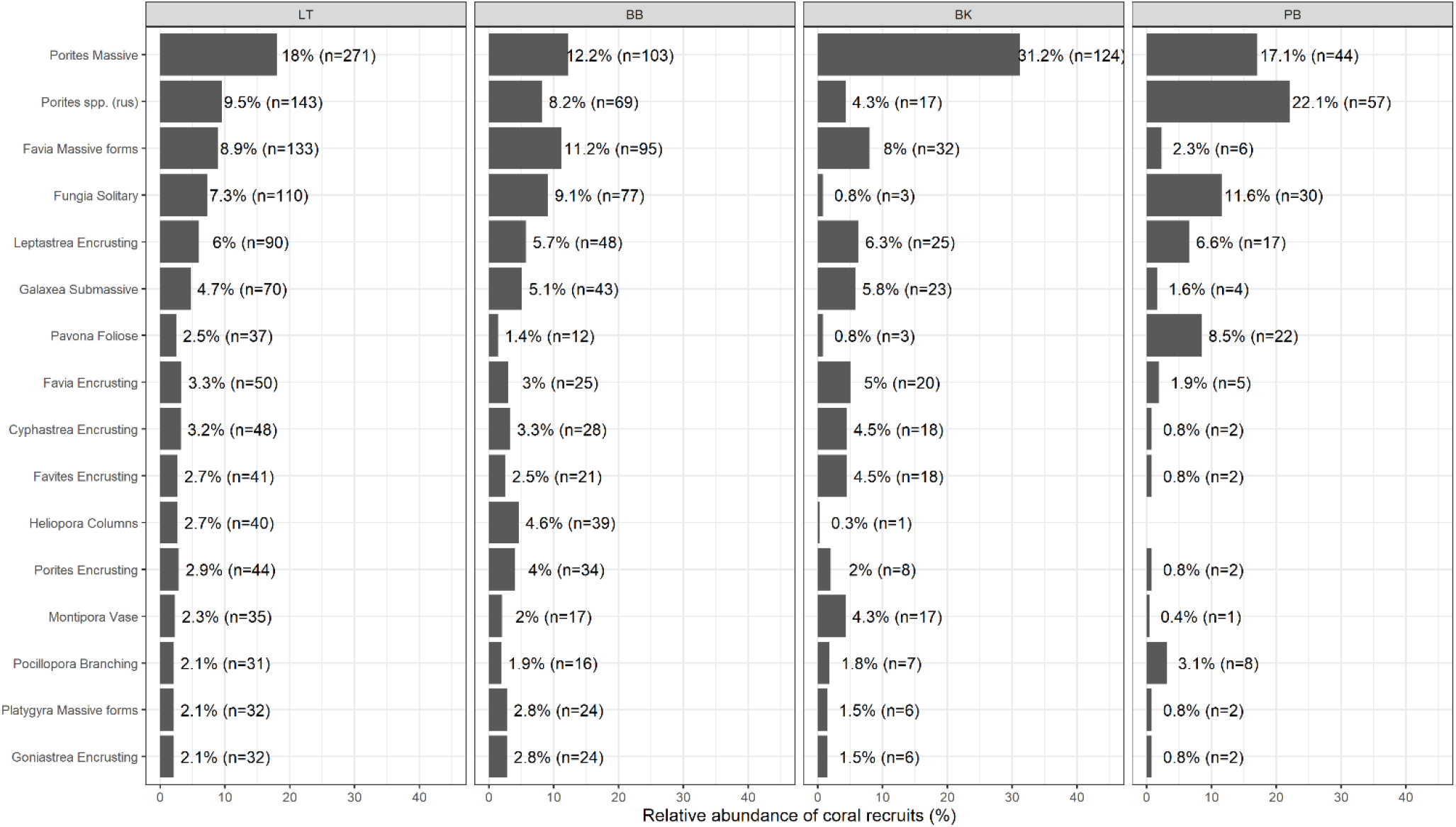
Percentage of relative coral recruitment is presented for the 16 most abundant hard coral morpho-taxa. The abundance of recruits (<5.0 cm) of each morpho-taxon is shown relative to the total number of recruits at regional (LT-Pulau Lang Tengah), and reef scale (BB-Batu Bulan; BK-Batu Kucing, PB-Pasir Besar).

The abundance of recruits relative to the respective abundance of adults varied considerably across sites for numerous morho-taxa (Supplementary S2).

### 3.3. Scleractinian size frequency distribution (SFD)

All log-transformed size-classes frequency distributions (SFDs) by morphology and site were non normal (SW tests p < 0.01). Similarly, most of SFDs by morpho-taxon and sites were non normal (74 % at BB (n=34), 81 % at BK (n=21), 87 % at PB (n=15). Scleractinian colony size was highly variable across and within sites. For instance, 273 of 561 (48.66%) moprho-taxonomic colony size comparisons at BB (Mann-Whitney-Wilcoxon test), 63 of 210 at BK (30.00% of pairs) and 67 of 105 (63.81% of pairs) were significantly different (electronic supplementary data).

Sample size was sufficient to test 37 morpho-taxa for differences in SFD across sites and within sites. Of these, 10 morpho-taxa were tested at all three sites, 13 at two sites and 14 at one site (Table 2, Supplementary Table S3), resulting in 70 morpho-taxonomic SFD tests across sites (n=3,799 colonies). Of the 70 morpho-taxonomic tests, SFD was positively skewed for 30 morpho-taxa, whereby only four were positively skewed at all sites *(encrusting Leptastrea,* massive *Lobophyllia,* encrusting *Pavona* and massive *Porites*) (Figure 4 and Figure 5). Of 39 negatively skewed size frequency distributions, eight were negatively skewed at all three sites: submassive *Galaxea*, *Porites spp (rus*), encrusting *Platygyra*, massive *Platygyra*, hispidose *Acropora*, laminar *Montipora*, solitary *Fungia* and massive *Symphyllia* (Table 2, Figure 4-6). Massive *Leptoria* was neither positively, nor negatively skewed (g_1_=0.00). Morpho-taxa tested at only one site are shown in the supplementary material Table S3.

**Figure 4.**
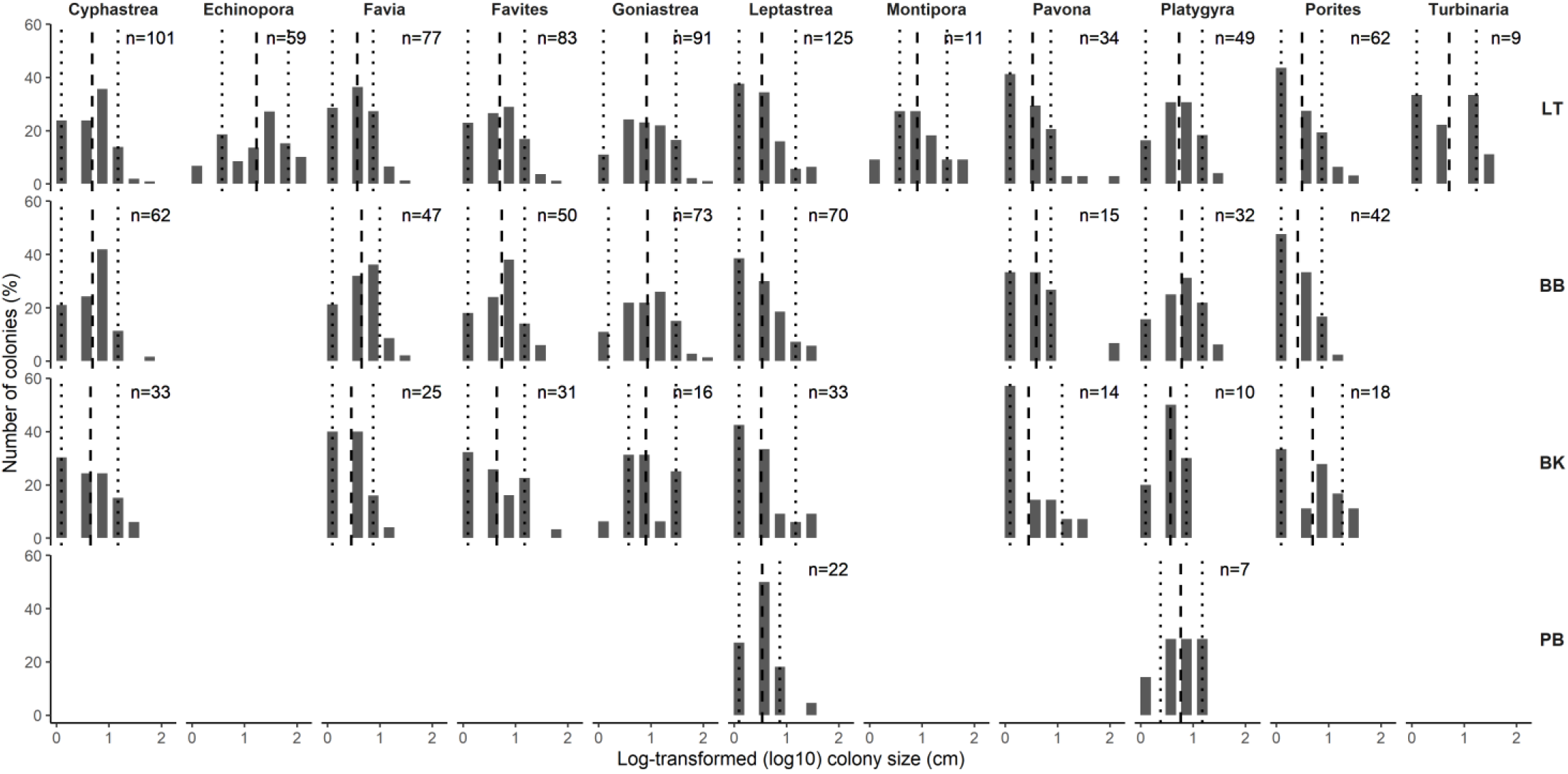
Log-transformed size frequency distribution (SFD) of encrusting hard coral morpho-taxa at reef scale at three sites (BB-Batu Bulan; BK-Batu Kucing; PB-Pasir Besar) and on island scale (LT-Lang Tengah), in Northeast Peninsular Malaysia. Dotted lines present the 10^th^ and 90^th^ percentile, respectively, and the dashed line shows the mean of the distribution.

**Figure 5.**
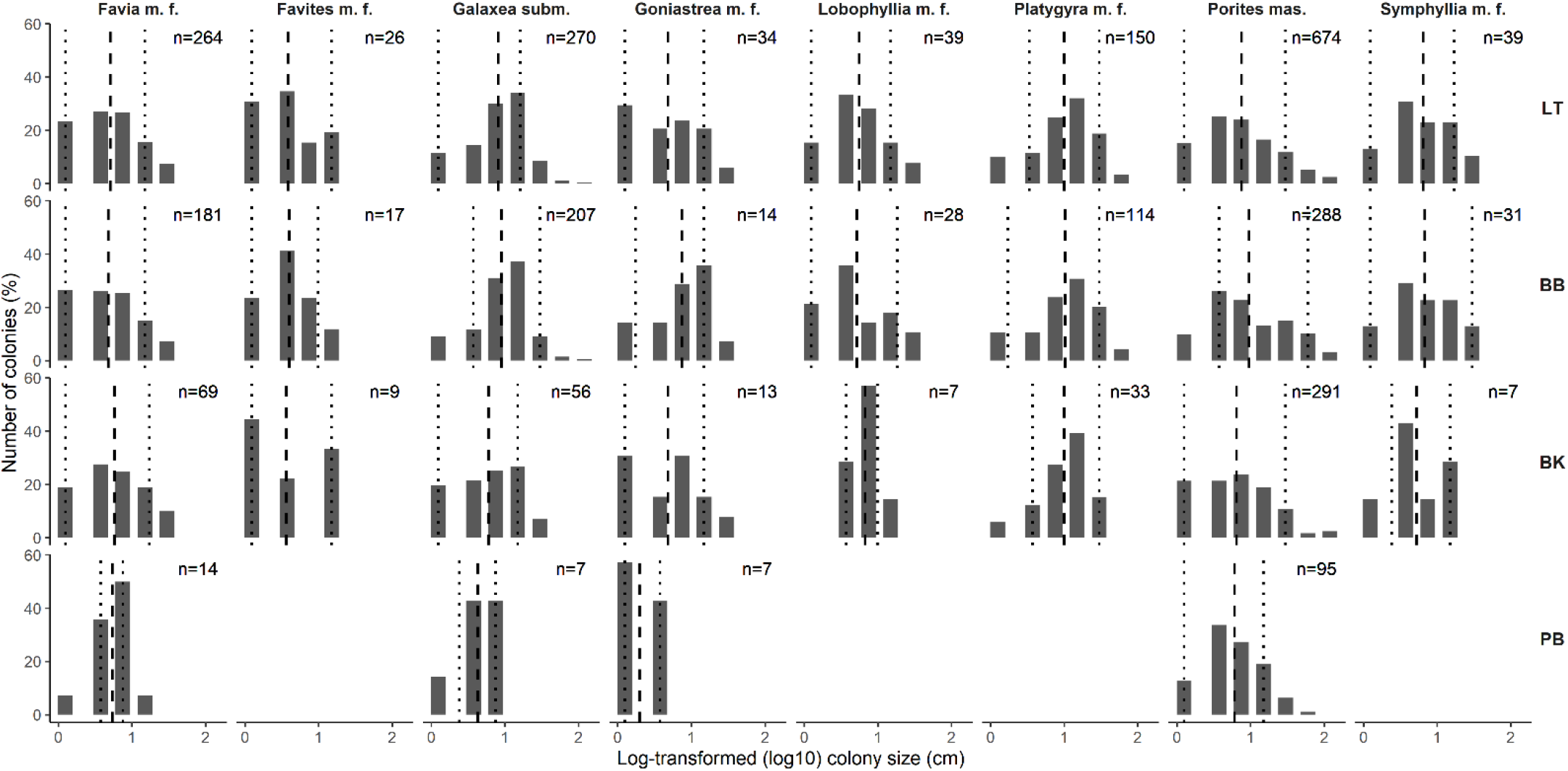
Log-transformed size frequency distribution (SFD) of massive hard coral morpho-taxa at reef scale at three sites (BB-Batu Bulan; BK-Batu Kucing; PB-Pasir Besar) and on island scale (LT-Lang Tengah), in Northeast Peninsular Malaysia. Dotted lines present the 10^th^ and 90^th^ percentile, respectively, and the dashed line shows the mean of the distribution. Abbreviations: m.f – massive forms (includes submassive and massive); subm.- submassive; mas.- massive.

**Figure 6.**
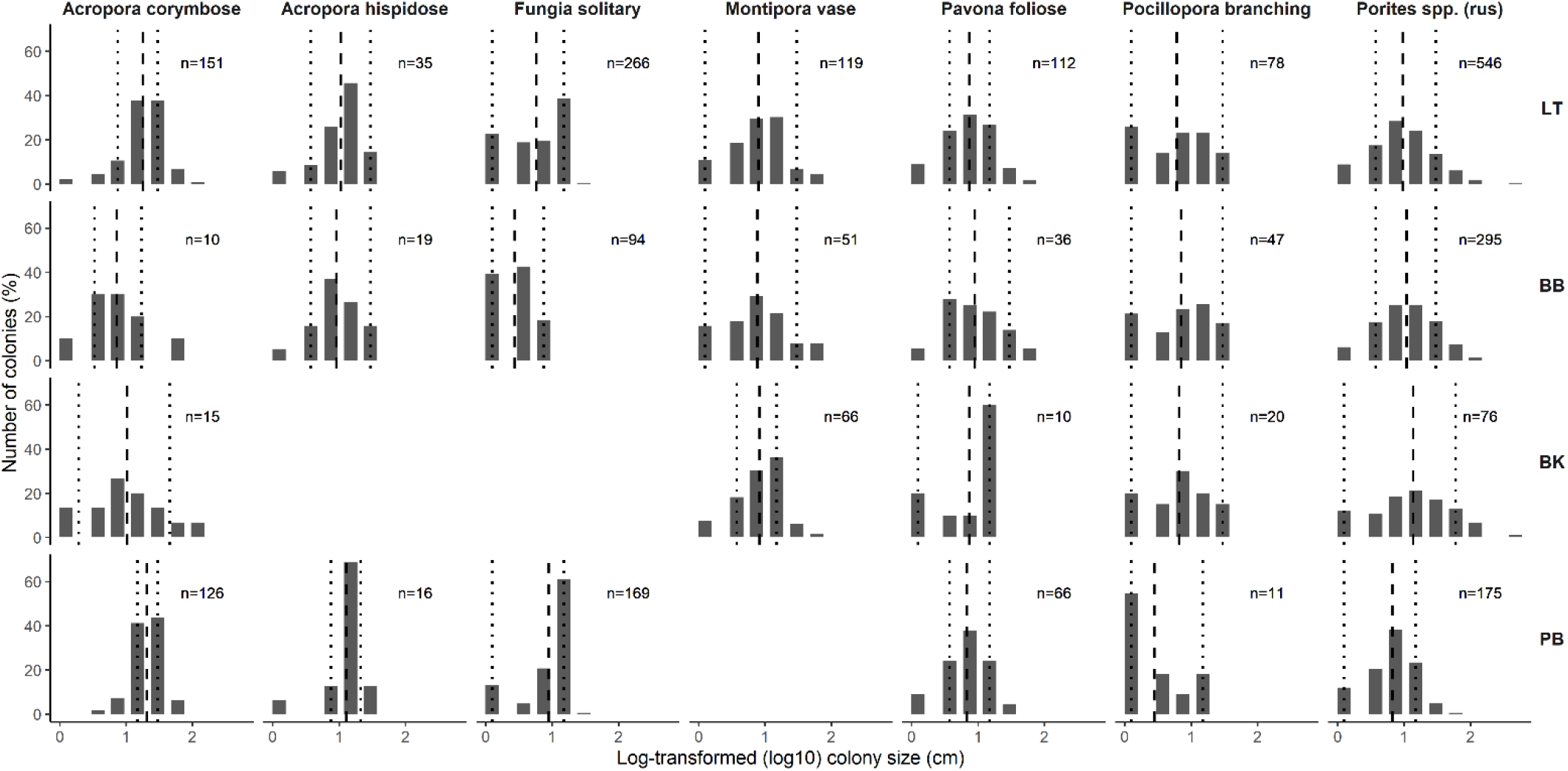
Log-transformed size frequency distribution (SFD) of hard coral morpho-taxa at reef scale at three sites (BB-Batu Bulan; BK-Batu Kucing; PB-Pasir Besar) and on island scale (LT-Lang Tengah), in Northeast Peninsular Malaysia. Dotted lines present the 10^th^ and 90^th^ percentile, respectively, and the dashed line shows the mean of the distribution.

**Table 2.** Size frequency distribution (SFD) metrics of scleractinian morpho-taxa at surveyed sites (BB-Batu Bulan; BK-Batu Kucing; PB-Pasir Besar) around Pulau Lang Tengah in Northeast Peninsular Malaysia. Anderson-Darling k-sample statistics show significant intra-taxonomic differences across sites. Highlighted in grey are the skewness and kurtosis distributions that are significantly different from the normal distribution as based on the Z-value > ±2.

| Morpho-taxon | Site | n | Size<sub>log</sub> (±SD) | 10<sup>th</sup> | 90<sup>th</sup> | g<sub>1</sub> (Z) | g<sub>2</sub> (Z) | CV % | AD k-test |
| --- | --- | --- | --- | --- | --- | --- | --- | --- | --- |
| *Acropora* corymbose | BB | 10 | 0.86 ± 0.46 | 0.53 | 1.24 | 0.42 (0.72) | 0.12 (0.16) | 53.18 | p < 0.001 |
| | BK | 15 | 1.01 ± 0.56 | 0.29 | 1.66 | 0.07 (0.14) | -0.48 (-0.61) | 55.22 | p < 0.001 |
| | PB | 126 | 1.31 ± 0.24 | 1.18 | 1.48 | -0.41 (-1.94) | 0.50 (1.22) | 18.01 | p < 0.001 |
| *Favia* massive | BB | 181 | 0.68 ± 0.43 | 0.10 | 1.18 | 0.05 (0.30) | **-0.98 (-2.81)** | 63.57 | ns |
| | BK | 69 | 0.76 ± 0.43 | 0.10 | 1.24 | -0.09 (-0.33) | -0.87 (-1.64) | 56.05 | ns |
| | PB | 14 | 0.73 ± 0.26 | 0.57 | 0.88 | -0.79 (-1.49) | 0.87 (1.12) | 35.24 | ns |
| *Galaxea* submassive | BB | 207 | 0.95 ± 0.38 | 0.57 | 1.48 | **-0.59 (-3.49)** | 0.51 (1.55) | 40.05 | p < 0.001 |
| | BK | 56 | 0.78 ± 0.43 | 0.10 | 1.18 | -0.32 (-1.02) | -0.93 (-1.63) | 54.75 | p < 0.001 |
| | PB | 7 | 0.63 ± 0.28 | 0.38 | 0.88 | -0.93 (-1.52) | -0.07 (-0.11) | 44.25 | p < 0.001 |
| *Goniastrea* massive | BB | 14 | 0.87 ± 0.41 | 0.24 | 1.18 | -0.71 (-1.33) | -0.38 (-0.48) | 47.29 | p ≤ 0.01 |
| | BK | 13 | 0.68 ± 0.47 | 0.10 | 1.18 | -0.03 (-0.05) | -1.18 (-1.51) | 69.07 | p ≤ 0.01 |
| | PB | 7 | 0.30 ± 0.26 | 0.10 | 0.57 | 0.29 (0.47) | **-1.92 (-2.9)** | 84.62 | p ≤ 0.01 |
| *Leptastrea* encrusting | BB | 70 | 0.54 ± 0.42 | 0.10 | 1.18 | 0.52 (1.86) | -0.65 (-1.23) | 78.18 | ns |
| | BK | 33 | 0.52 ± 0.45 | 0.10 | 1.18 | **0.80 (2.05)** | -0.36 (-0.53) | 87.25 | ns |
| | PB | 22 | 0.54 ± 0.35 | 0.10 | 0.88 | 0.53 (1.17) | 0.66 (0.87) | 64.16 | ns |
| *Pavona* foliose | BB | 36 | 0.95 ± 0.42 | 0.57 | 1.48 | 0.08 (0.22) | -0.47 (-0.70) | 43.96 | ns |
| | BK | 10 | 0.87 ± 0.45 | 0.10 | 1.18 | -0.98 (-1.69) | -0.72 (-0.96) | 52.05 | ns |
| | PB | 66 | 0.83 ± 0.34 | 0.57 | 1.18 | -0.47 (-1.64) | 0.01 (0.02) | 40.83 | ns |
| *Platygyra* encrusting | BB | 32 | 0.78 ± 0.39 | 0.10 | 1.18 | -0.30 (-0.76) | -0.60 (-0.87) | 50.51 | ns |
| | BK | 10 | 0.57 ± 0.28 | 0.10 | 0.88 | -0.63 (-1.09) | -0.62 (-0.82) | 49.95 | ns |
| | PB | 7 | 0.76 ± 0.38 | 0.38 | 1.18 | -0.54 (-0.87) | -0.67 (-1.01) | 50.18 | ns |
| *Pocillopora* | BB | 47 | 0.85 ± 0.48 | 0.10 | 1.48 | -0.40 (-1.2) | -1.05 (-1.72) | 56.33 | p < 0.05 |
| | BK | 20 | 0.82 ± 0.46 | 0.10 | 1.48 | -0.31 (-0.67) | -0.92 (-1.21) | 56.28 | p < 0.05 |
| | PB | 11 | 0.45 ± 0.45 | 0.10 | 1.18 | 0.67 (1.18) | -1.15 (-1.50) | 99.59 | p < 0.05 |
| *Porites* spp. (rus) | BB | 295 | 1.04 ± 0.44 | 0.57 | 1.48 | -0.12 (-0.82) | -0.26 (-0.95) | 42.20 | p < 0.001 |
| | BK | 76 | 1.14 ± 0.59 | 0.10 | 1.78 | -0.07 (-0.24) | -0.36 (-0.71) | 51.37 | p < 0.001 |
| | PB | 175 | 0.83 ± 0.37 | 0.10 | 1.18 | **-0.43 (-2.34)** | -0.07 (-0.19) | 44.33 | p < 0.001 |
| *Porites* massive | BB | 288 | 0.98 ± 0.52 | 0.57 | 1.78 | 0.21 (1.44) | **-0.69 (-2.47)** | 52.68 | p < 0.001 |
| | BK | 291 | 0.81 ± 0.50 | 0.10 | 1.48 | 0.20 (1.43) | -0.45 (-1.62) | 61.64 | p < 0.001 |
| | PB | 95 | 0.78 ± 0.38 | 0.10 | 1.18 | 0.01 (0.05) | -0.29 (-0.63) | 49.35 | p < 0.001 |
| *Acropora* hispidose | BB | 19 | 0.96 ± 0.36 | 0.57 | 1.48 | -0.46 (-0.95) | 0.11 (0.14) | 37.02 | ns |
| | PB | 16 | 1.11 ± 0.31 | 0.88 | 1.33 | **-2.18 (-4.28)** | **5.25 (6.75)** | 28.09 | ns |
| *Cyphastrea* encrusting | BB | 62 | 0.69 ± 0.37 | 0.10 | 1.18 | -0.13 (-0.45) | -0.06 (-0.11) | 54.30 | ns |
| | BK | 33 | 0.65 ± 0.44 | 0.10 | 1.18 | 0.10 (0.27) | -1.08 (-1.59) | 68.23 | ns |
| *Favia* encrusting | BB | 47 | 0.65 ± 0.36 | 0.10 | 1.00 | -0.19 (-0.57) | -0.54 (-0.89) | 54.46 | p ≤ 0.01 |
| | BK | 25 | 0.46 ± 0.33 | 0.10 | 0.88 | 0.26 (0.60) | -1.00 (-1.37) | 72.96 | p ≤ 0.01 |
| *Favites* encrusting | BB | 50 | 0.74 ± 0.39 | 0.10 | 1.18 | -0.23 (-0.70) | -0.51 (-0.86) | 52.30 | ns |
| | BK | 31 | 0.64 ± 0.47 | 0.10 | 1.18 | 0.31 (0.78) | -0.75 (-1.08) | 72.71 | ns |
| *Favites* massive | BB | 17 | 0.60 ± 0.35 | 0.10 | 1.00 | -0.09 (-0.19) | -0.84 (-1.08) | 58.35 | ns |
| | BK | 9 | 0.56 ± 0.50 | 0.10 | 1.18 | 0.31 (0.52) | **-1.62 (-2.20)** | 88.79 | ns |
| *Fungia* solitary | BB | 94 | 0.44 ± 0.30 | 0.10 | 0.88 | -0.01 (-0.02) | **-1.45 (-3.10)** | 67.76 | p < 0.001 |
| | PB | 169 | 0.95 ± 0.37 | 0.10 | 1.18 | **-1.49 (-8.04)** | **0.87 (2.41)** | 38.99 | p < 0.001 |
| *Goniastrea* encrusting | BB | 73 | 0.93 ± 0.45 | 0.19 | 1.48 | -0.11 (-0.39) | -0.36 (-0.69) | 48.58 | ns |
| | BK | 16 | 0.90 ± 0.42 | 0.57 | 1.48 | 0.08 (0.16) | -0.80 (-1.03) | 46.06 | ns |
| *Lobophyllia* massive | BB | 28 | 0.72 ± 0.45 | 0.10 | 1.27 | 0.14 (0.33) | -0.97 (-1.36) | 62.18 | ns |
|  | BK | 7 | 0.83 ± 0.21 | 0.57 | 1.00 | 0.13 (0.22) | -0.61 (-0.92) | 24.97 | ns |
| *Montipora* laminar | BB | 51 | 0.88 ± 0.48 | 0.10 | 1.48 | -0.03 (-0.09) | -0.52 (-0.87) | 54.00 | ns |
|  | BK | 66 | 0.92 ± 0.36 | 0.57 | 1.18 | -0.53 (-1.83) | 0.27 (0.49) | 39.07 | ns |
| *Pavona* encrusting | BB | 15 | 0.6 ± 0.52 | 0.10 | 0.88 | **1.45 (2.79)** | **2.50 (3.2)** | 87.00 | ns |
|  | BK | 14 | 0.45 ± 0.48 | 0.10 | 1.09 | 0.92 (1.72) | -0.51 (-0.65) | 105.8 | ns |
| *Platygyra* massive | BB | 114 | 1.01 ± 0.44 | 0.24 | 1.48 | **-0.61 (-2.72)** | -0.16 (-0.36) | 43.18 | ns |
|  | BK | 33 | 1.00 ± 0.36 | 0.57 | 1.48 | **-0.85 (-2.18)** | 0.52 (0.77) | 35.48 | ns |
| *Porites* encrusting | BB | 42 | 0.41 ± 0.33 | 0.10 | 0.88 | 0.40 (1.15) | -1.14 (-1.80) | 79.98 | p < 0.001 |
|  | BK | 18 | 0.70 ± 0.50 | 0.10 | 1.27 | -0.01 (-0.02) | -1.33 (-1.72) | 71.54 | p < 0.001 |
| *Symphylia* massive | BB | 31 | 0.83 ± 0.42 | 0.10 | 1.48 | -0.18 (-0.46) | -0.76 (-1.10) | 50.31 | ns |
|  | BK | 7 | 0.72 ± 0.39 | 0.38 | 1.18 | -0.19 (-0.31) | -0.89 (-1.35) | 53.49 | ns |

SFD of the following 11 morpho-taxa varied across sites: corymbose *Acropora,* encrusting *Cyphastrea,* encrusting *Favia,* massive *Favia,* encrusting *Favites,* massive *Favites,* encrusting *Goniastrea,* massive *Goniastrea,* foliose *Pavona, Pocillopora,* encrusting *Porites* (Table 2). Here, there were significant differences across sites for five morpho-taxa with varying SFD across sites: corymbose *Acropora* (AD k-test p<0.001), massive *Goniastrea* (AD k-test p≤0.01) and *Pocillopora* (AD k-test p<0.05), encrusting *Porites*, (AD k-test p<0.001), and encrusting *Favia*. Furthermore, significant differences in SFD were found for taxa whose SFD was negatively skewed at all three sites, such as submassive *Galaxea* (AD k-test p<0.001), *Porites spp. (rus)* (AD k-test p<0.001) and solitary *Fungia* (AD k-test p<0.001). Of morpho-taxa with positive SFD at all three sites, massive *Porites* showed significant differences in size distribution across sites (AD k-test p<0.001). Overall, leeward PB was more dominated by negatively skewed morpho-taxa (11 of 15, 73.33%) than BK (11 of 21, 52.38%) and BB (18 of 34, 52.94%). Similarly, leeward PB was more leptokurtic compared to BB and BK (Table 1): PB 40.00% leptokurtic morpho-taxa, 9.52% at BK, and 17.65% at BB. In addition to morpho-taxonomic SFD differences across sites, there were further differences in morpho-taxa SFD within sites, with 28.16% (158 of 561) of morpho-taxonomic SFD pairs being significantly different at BB, 10.47% (22 of 210) at BK and 49.52% (52 of 105) at PB (electronic supplementary data).

The Z-statistics of the SE Skewness of all 70 tested morpho-taxa revealed a significantly different SFD from the normal distribution for eight morpho-taxa (Z>±2), and the Z-statistics of the SE kurtosis highlighted significantly different SFD (Z>±2) for nine morpho-taxa (Table 2). Noteworthy, hispidose *Acropora* at PB, solitary *Fungia* at PB and encrusting *Pavona* at BB (g_1_=1.45, g_2_=2.50, CV=87.00%) were both, significantly skewed and peaked. Here, hispidose *Acropora* and solitary *Fungia* were significantly skewed towards larger colonies (g_1_= −2.18 and −1.49, respectively), colony size was highly centralized (leptokurtic, g_2_=5.25 and 0.87, respectively), and less diverse (CV=28.09% and CV=38.99%, respectively). In contrast to PB, kurtosis of solitary *Fungia* at BB was negative (platykurtic; g_2_= −1.45, CV=67.76%). At BB, all nine massive morpho-taxa were platykurtic, whereby massive *Favia* (g_2_= −0.98, CV=63.57%) and massive *Porites* (g_2_= −0.69, CV=52.68%) were significantly platykurtic (Table 2). At BK, kurtosis was significant for massive *Favites* (g_2_= −1.62, CV=88.79%). In addition to encrusting *Pavona*, encrusting *Turbinaria* at BB was the only other encrusting morpho-taxa with significant kurtosis (g_2_= −1.48, CV= 84.48%) (Supplementary S3). Skewness was significantly negative for massive *Platygyra* at BB (g_1_= −0.61) and BK (g_1_= −0.85), *Porites spp. (rus)* at PB (g_1_= −0.43) and submassive *Galaxea* at BB (g_1_= −0.59). Encrusting *Leptastrea* was significantly positively skewed at BK (g_1_=0.80).

## 4. Discussion

### 4.1. Benthic surveys and reef health

Benthic surveys showed marked and consequential differences in reef health based on benthic substrate composition and general reef health indicators (Table 1). All indicators (e.g., taxonomic diversity and richness, colony density, hard coral cover, and colony size CV), were lower at leeward PB and higher at windward BB. Live hard coral cover was markedly low at leeward PB (8.78%), where coastal development possibly resulted in physical reef degradation (Figure 1). In addition, herbivorous fish abundance is reportedly lower at PB than BK and BB (CK. Lynn, unpublished data, 2018). Conceivably, secondary impacts from low abundance of herbivorous fish (McClanahan et al., 2001; Green and Bellwood, 2009), sewage discharge and sedimentation rates may persistently impact coral reef health at leeward sites (Wooldridge et al., 2012; Duckworth et al., 2017; Reef Check Malaysia, 2019), resulting in lower hard coral colony diversity, abundance and less variable colony size at leeward PB (Table 1). Although historical data is not available and temporal analysis of percent live hard coral cover is not possible, the disproportionately high amount of coral rubble at PB (39.61 %) compared to BB (11.82 %), and the declining trend of coral reef health and cover across Peninsular Malaysia (Toda et al., 2007; Praveena et al., 2012; Reef Check Malaysia, 2019), suggests that much of the observed differences in benthic substrate composition and reef health indicators are substantially driven by human activity. As such, coral reefs in close proximity to coastal developments are particularly impacted, and a clear gradient of reef health is evident and subsequent differences in coral assemblages are consequential (Table 1 and Figure 2).

### 4.2. Demographic structure and recruitment

Coral recruitment was dominated by a few morpho-taxa, such as massive *Porites, Porites spp. (rus),* massive *Favia* and solitary *Fungia* (Figure 3, Supplementary S2 and S4). According to findings by Riegl et al. (2012), constant recruitment impulses and declines in colony size after mass disturbance events underpin demographic structure, and McClanahan et al. (2001; 2008) further demonstrated that SFD significantly shifts to smaller colonies after mass ecological disturbance (e.g., coral bleaching) and due to chronic stress (e.g., overfishing). Accordingly, chronically disturbed reef environments should host a larger amount of recruits and sub-adult colonies, but may also be constrained by high post-settlement mortality (Chong-Seng et al., 2014). Pulau Lang Tengah’s reefs are chronically stressed by human impacts, and during this study we identified a generally high number of coral recruits (e.g., 36.94% of the surveyed population), and smaller sized sub-adult colonies (e.g., < 20.00 cm; 47.08% of the surveyed population). Moreover, reef scale assessments further revealed that healthier and more intact sites (here BB) were hosting a higher abundance of both, larger (adult) and smaller colonies (sub-adult and recruits), as compared to more degraded sites such as BK and PB (Table 1). However, the general demographic structure was consistent across sites, as smaller coral colonies of 10-20 cm in size were the most preponderant (Table 1). Presumably, this preponderance of smaller colonies may be the result of extrinsic demographic factors such as large-scale disturbance events that eliminated larger colonies and induced shifts in the abundance and dominance of individual taxa (Meesters et al., 1996; Bak and Meesters, 1998, 1999). Correspondingly, after large-scale disturbance events, such as the 1998 and 2010 mass bleaching events (Kushairi, 1999; Tan and Heron, 2011), morpho-taxa such as massive *Porites* have started to replace formerly dominant taxa such as *Acropora* and *Montipora* (Brown, 1997; Toda et al., 2007). Preconditions for such taxon-specific shifts would be visible in the taxon’s current SFD and would be specifically highlighted by a platykurtic distribution. Furthermore, in view of skeletal growth rates of 2.0 cm / year of massive *Porites* colonies measured in nearby (∼10 km) Redang Island (Tanzil et al., 2013), and considering the time elapsed since the last mass coral bleaching events with significant mortality rates (nine and 21 years, respectively), impulses of post-disturbance recruitment are likely to have contributed to the preponderance of smaller size groups and the co-aligned dominance of a few taxa, such as massive *Porites*, and *Porites spp. (rus).* (Table 1, Supplementary S4). Such findings are similar to studies from the Red Sea and French Polynesia (Riegl et al., 2012; Adjeroud et al., 2015), where the demographic structure was dominated by recruits and small colonies on reefs that suffered recent and frequent ecological disturbances. On the other hand, frequent disturbances and chronic stressors may have induced a decline in general colony size. For instance, in Kiribati, chronic and multiple stressors resulted in a decadal and general decline in colony size of the six most dominant taxa, declining to an average size of between 10-20 cm, (Cannon et al., 2021), and further resulted in the dominance of taxa such as *Porites spp. (rus)*. These observations are supported by contrasting colony sizes across gradients of reef health, as all major taxa (except for corymbose *Acropora* and solitary *Fungia*), were larger at healthier windward BB than at PB (Table 2). Nonetheless, pre-existing conditions specific to each reef site, and conditions resulting from ecological disturbances (e.g., substrate suitability and availability), are likely further determining recovery on reef-scale, and are key in understanding the demographic structure and resilience of hard corals on individual reefs.

For significant shifts in the mix of taxa and SFD to occur after mass disturbance due to successful coral recruitment, ample and suitable substrate is necessary to prevent high post-settlement mortality (Chong-Seng et al., 2014). If unavailable, selective pressures may result in recruitment bottlenecks in favour of fewer taxa that are capable of successful recruitment in unfavourable environments, such as rubble dominated reefs. We investigated successful coral recruitment by quantifying the abundance of coral recruits (Table 1, Figure 3), in view of site-specific benthic substrate composition. Our results suggest that coral recruitment abundance and diversity was consistent with patterns of site-specific coral reef health. As such, coral recruitment was lowest at PB and highest at BB (Table 1, Supplementary S2). Presumably, the bottom-substrate composition of the reef mosaic strongly impacts coral recruitment at reef scale (Chong-Seng et al., 2014). For instance, at PB, coral rubble and sand where the most dominating substrates (Table 1). Coral rubble in particular, is tagged as a coral recruitment inhibitor and thus prevents the establishment of mature colonies, resulting in demographically unviable populations (Raymundo et al., 2007; Wolfe et al., 2021), and in the preponderance of a few taxa, that are able to reproduce and settle in such environments, such as solitary *Fungia*, foliose *Pavona*, massive *Porites* and *Porites spp. (rus)*. Conversely, studies from the tropical Pacific suggest that the recruitment abundance of *Acropora*, *Pocillopora* and *Porites* are correlated with the abundance of adults. (Penin and Adjeroud, 2013; Bramanti and Edmunds, 2016). This may partially explain why dominant adult morpho-taxa are further dominating the recruitment pool, and recruitment dominance of individual taxa such as *Acropora*, *Porites* and *Fungia* was noted in Peninsular Malaysia on artificial reefs and on *in-situ* tiles (Rani et al., 2015; Hanapiah et al., 2017). Lastly, species-specific recruitment mortality rates may further explain the dominance of a fewmorpho-taxa (Smith, 1992).

### 4.3. Scleractinian colony size frequency distribution across reef health

It is important to note that historical records of hard coral SFD are unavailable for Peninsular Malaysia and therefore retrospective conclusions about disturbance induced shifts in SFD are not feasible, neither is it possible to retrospectively ascertain the importance of disturbance events on the present demographic SFD. This study intended to create a baseline reference for future studies, but it must be noted that changes in demographic structure are subject to strong spatio-temporal patterns and variability (particularly coral recruitment), and conclusions are only momentarily. Further and repeated studies on species-specific level are needed to quantify the impacts of global and local stressors on demographic structure in order to highlight the subsequent changes in community assemblages of hard corals (Edmunds and Riegl, 2020).

To determine present variations in hard coral demography and population viability in view of chronic disturbances and human stressors, it was necessary to compare SFD across reefs in various stages of reef health and across reefs subjected to the same overarching stressors. We compared hard coral SFD across three reef sites, which were in good (BB > 40% hard coral cover), fair (BK > 20% hard coral cover) and degraded conditions (PB < 20% hard coral cover) (Table 1). SFD of 11 morpho-taxa was varied across sites, eight morpho-taxa were negatively skewed at all three sites, and four taxa were positively skewed at all sites (Table 2, Figures 4-6). We found evidence for both proposed theories of shifts to either large or small colonies. For instance, on island scales, this study recorded a preponderance of small to medium sized corals (Table 1), which possibly resulted from a recent demographic disturbance event followed by recruitment (e.g., massive *Porites* as described above), and could further reflect the consequences of chronic and multiple stress, similar to reefs in Kiribati (Cannon et al., 2021). Variations in SFD were highly heterogenic across morpho-taxa, however, we found pronounced differences with morpho-taxa across reef health (Table 2, Figures 4-6), which offer insights into the impacts of various site stressors on taxon-specific SFD (Bauman et al., 2013). Overall, the most degraded site (PB) was dominated by colony SFD shifted to larger colonies, a more leptokurtic distribution, low recruitment density, as well as generally lower hard coral abundance and diversity (Table 1-2, Figure 3). Moreover, the taxonomic assemblage here was dominated by weedy taxa (Supplementary S5). These findings thus partially confirm postulates by Meesters et al. (2001) and Dietzel et al. (2020), who argued that degraded reefs are characterized by large colonies and low colony size variability, leading to a depression in recruitment and subsequent reef replenishment. Nevertheless, whereby our results highlight the limited demographic viability of more disturbed sites (Table 2, Figures 4-6), results further revealed differences in SFD of numerous morpho-taxa across sites. As such, some morpho-taxa were larger at healthier sites (here BB) compared to degraded sites. For instance, whereas submassive *Galaxea* was negatively skewed at all sites, skewness was only significant at BB, where it attained its largest size on average (Table 2). Massive *Platygyra* was significantly skewed towards larger colonies at healthier sites, and both taxa were abundant as recruits (Figure 3). SFD of encrusting taxa such as encrusting *Favia* were significantly different at BB (good condition and negative distribution) compared to BK (fair condition and positive distribution). Thus, negative skewness is not in immediate association with reef degradation (*sensu* Meesters et al., 2001; Miller et al., 2016; Dietzel et al., 2020), but is taxon-dependent and likely due to the inherent life-history strategies of present taxa. Possibly, resistance to environmental stress, such as heat stress and subsequent bleaching, favours negative skewness and larger size of taxa that are more tolerant to such conditions (Bauman et al., 2013). Submassive *Galaxea* is locally a more bleaching tolerant taxon (Szereday and Affendi, 2022), and possibly attains larger size on reefs with intact framework and sufficient substrate (such as BB), whilst taxa susceptible to heat stress are reduced in size and are filtered out (McClanahan et al., 2001, 2008), offering vacant substrate for growth and reducing spatial competition.

Contrasting the indicators of reef health against colony SFD, it is plausible that coral reef degradation results in a SFD tipping point, where fewer morpho-taxa are able to maintain viable demographic populations as coral reef degradation deteriorates the reef substrate and structure, and ultimately exacerbates the loss of colonies. Such factors are also possible drivers of phase shifts, where complex reef assemblages are replaced by rudimentary and weedy taxa (McWilliam et al., 2020). Indeed, site-specific variations of hard coral SFD are due to the reef-specific presence of chronic stressors, as well as life-history traits, that enable certain coral taxa to persist under chronic and multiple stress (Darling et al., 2012, 2013). Chronic disturbances and mortality shocks would result in a leptokurtic distribution of impacted taxa at disturbed sites, and generally in platykurtic distribution of tolerant taxa, especially at healthier sites (Bauman et al., 2013). Weedy morpho-taxa such as solitary *Fungia* and foliose *Pavona* dominated the assemblage on the most degraded reef site (PB) (Figure 2), in addition to corymbose and hispidose *Acropora,* as well as *Porites spp. (rus).* Moreover, high population turnover is characterized by leptokurtic distribution and is associated with frequent disturbances (Kayal et al., 2015), noted for these taxa, with the exception of *Porites spp.*(*rus*). In contrast, platykurtic distribution was predominantly characteristic of massive taxa, particularly at less degraded sites, and was significant for massive *Favia*, solitary *Fungia*, massive *Porites* at BB, and massive *Favites* at BK (Table 2).

The current state of coral reef health and the biophysical interaction of monsoon waves with the predominant reef substrates (e.g., coral rubble, live hard coral), provides a diagnostic glimpse into the consequences of severe reef degradation, and offers further explanation of demographic tipping points of individual reef sites. In this realm, natural recovery is significantly impeded due to biophysical interactions, as in Pulau Lang Tengah specifically, the northeast monsoon winds induce strong wind and wave action (Figure 1B), which establishes a system in a state of hysteresis (*sensu* Hughes et al., 2010), where the excessive amount of coral rubble, sand and monsoon waves interact to trigger further decline in hard coral colony abundance, due to sedimentation and physical fragmentation. This state of hysteresis hypothetically prevents successful larvae settlement and development, as suggested by the low rates of recruitment and the high substrate cover of rubble and sand at PB. Thus, the combination of reinforcing factors increases colony mortality and possibly further reduces the local larvae source pool. Secondly, such reinforcing feedbacks increases the amount of mobile substrate that acts as the degenerating agent. Therefore, once reef substrate degradation reaches a critical level at which the proposed reinforcing mechanisms interact in a strong feedback loop, demographic recovery is reduced and only taxa with beneficial life-history strategies and physical properties with high hydro-mechanical tolerance, are able to persist. For instance, hispidose *Acropora* (e.g., *Acropora longicyathus*), is known to persist well in coral rubble fields (Veron et al., 2022), which may be due to colony morphology as well as life-history characteristics, such as quick growth, density dependent recruitment and asexual reproduction by fragmentation. Indeed, Szereday and Affendi confirmed a high dominance of this taxon at PB during repeated annual surveys (Szereday and Affendi, 2022). Secondly, solitary *Fungia* was particularly dominant in Lang Tengah’s coral rubble fields, likely due to the non-sessile lifestyle and colony shape that enables survival in shallow turbulent waters (Jokiel and Cowdin, 1976), as well as secondary life-history strategies (*sensu* Darling et al., 2013), which imply that this taxon can benefit from degraded and unproductive environments. Thus, SFD of solitary *Fungia* was significantly different between BB and PB (Table 2). At PB the dominance of larger *Fungia* colonies was underlined by leptokurtic distribution and negative skewness, in sharp contrast to BB. Other negatively skewed taxa at PB were corymbose *Acropora* and *Porites spp. (rus)*. SFD of *Porites spp. (rus)* at PB contrasted BB and BK, as at healthier sites it was more evenly distributed and thus demographically more viable. Although this taxon occurs in very dense coral carpets and visibly contributes to the local leeward reef frame building, high physical fragmentation was observed at PB, which is likely a result of human disturbances (e.g., coastal development, sedimentation), and subsequently weakens the reef frame and magnifies wave induced sedimentation. Furthermore, in the case of corymbose *Acropora,* the lack of small colonies as well as low coral recruitment numbers may be due to unsuitable substrate. With increasing degradation and annually occurring monsoon waves, fragmentation possibly increases due to wave action and collision with mobile rubble. Indeed, studies suggest that such broken fragments have limited reattachment and survival success in rubble fields (Cameron et al., 2016). Occurrence in high density and mono-specific coral carpets may provide enough robustness to withstand monsoon waves, but the lack of suitable substrate would impede reef-wide larvae dispersal (Chong-Seng et al., 2014), which would be otherwise warranted due to high adult abundance (*sensu* Bramanti and Edmunds, 2016). Equally, asexual reproduction is hindered as reattachment of broken coral fragments is directly influenced by fragment size and the benthic substrate composition (Smith & Hughes, 1999). Therefore, it is possible that only larger fragments of certain growth-from (e.g., hispidose branching) can quickly reattach to the reef matrix, causing lower colony size range and result in the observed leptokurtic distribution. Ultimately, in environments dominated by an annual monsoon and mobile substrates, the present demographic structure of these taxa represents a severe reproductive barrier, threatening long-term population viability (Kramer et al., 2021). This may also partially explain why only four taxa (corymbose *Acropora*, solitary *Fungia*, massive *Porites* and *Porites spp. (rus)*) dominated ∼70% of the assemblage at PB, in contrast to BB, where 11 taxa constituted ∼70% of the assemblage (Supplementary S5), while SFD was significantly more negative here than compared to less degraded sites (Table 1). Conclusively, such reefs are compromised in their capacity to suppress the movement and resuspension of sand and rubble, reinforcing hysteresis between sedimentation and monsoon waves, resulting in the survival of only larger colonies in dense clusters (e.g., corymbose *Acropora*, foliose *Pavona*, *Porites spp. (rus)*), and with leptokurtic distribution (e.g., solitary *Fungia*, corymbose and hispidose *Acropora*).

Despite indication that SFD can re-balance after mass disturbance such as hurricanes and coral bleaching (Crabbe, 2009), the presented system of hysteresis and the complex interaction of biophysical drivers would prevent significant recovery and result in depleted rubble environments, as well as in reef assemblages with significant deficits in functional trait diversity (McWilliam et al., 2021), leading to a loss of reef complexity and diversity, regardless of whether SFD shifts towards a negatively or positively distributed SFD. Substrate stabilization has shown to be a viable tool to increase hard coral cover (Fox et al., 2019), but scalability remains an issue (Williams et al., 2018), and with continuous anthropogenic heating (Szereday and Affendi, 2022), such management tools may soon become unviable. Nonetheless, several taxa showed high absolute abundance, high recruitment rates and an evenly distributed demographic structure, which is further coherent with findings from other regions (e.g., French Polynesia), highlighting taxa that might be able to persist under continuous stress and environmental degradation. For instance, while massive colonies were generally smaller at PB compared to BB and BK, no clear differences in distribution metrics (colony size CV, skewness and kurtosis) were discernible across sites for massive taxa (Supplementary S6), which suggests general resistance to prevailing conditions at all sites and less selective pressure on colony size and SFD. In French Polynesia, massive *Porites* has been shown to be more resistance to perturbations due to size-structure homogeneity (Adjeroud et al., 2007, 2015), typically due to low individual turnover, high longevity and resistance to environmental perturbations (Kayal et al., 2015). Here, characteristics of massive *Porites* were high recruitment rates and a stable demographic structure (as indicated by insignificant skewness and significant platykurtic distribution), similar to the central Pacific (Adjeroud et al., 2015). This was further true for several massive taxa (e.g., *Favia*, *Porites*, *Platygyra* and *Goniastrea*), whereby dominance and steadiness of massive taxa can be explained by longevity and fecundity (Alvarez-Noriega et al., 2016), as well as hydrodynamic stability (Madin et al., 2014). In Kiribati, *Porites spp. (rus)* increased in relative abundance after multiple disturbances to dominate the reef assemblage (Cannon et al., 2021), emerging as a more tolerant and stress resistant taxon. Indeed, *Porites spp. (rus)* colonies have shown acclimation potential to turbid and nutrient-rich environments by diversifying strategies for energy acquisition to facilitate persistence (Padilla-Gamiño et al., 2012). Considering the high levels of sedimentation and the probable sewage discharge by nearby resorts at PB, such energy acquisition strategies may co-explain the persistence of *Porites spp. (rus)* at PB. Ultimately, dominant morpho-taxa (e.g., massive *Favia*, columnar *Heliopora*, massive *Porites*, *Porites spp. (rus)*) displayed high levels of recruitment, absolute abundance, platykurtic distribution and were neither negatively nor positively skewed at significant levels (Table 2). This indicates population stability and low individual turnover rates, which further highlight demographic viability and tolerance to perturbations. However, only the two *Porites* taxa were able to persist in significant numbers at the severely degraded site, highlighting the selective pressure of the proposed tipping point.

## 5. Conclusion

In conclusion, healthier sites were richer in hard coral cover, recruitment, abundance and diversity (Table 1). Chronic and widespread stress represents selective pressures on successful recruitment, in favour of a limited amount of morpho-taxa that survive disturbance as adults and reproduce successfully (e.g., massive *Porites, Porites spp. (rus)*), or are morpho-taxa that are generalist, cryptic or weedy (e.g., solitary *Fungia*, encrusting *Leptastrea*, foliose *Pavona*). Ultimately, such assemblages are significantly different compared to healthier and more intact reef sites (e.g., PB vs BB) (Figure 2). Under multiple disturbance regimes and chronic stress, a sharp decline in the abundance of all size classes is likely (Dietzel et al., 2020; Cannon et al., 2021), in addition to a shift towards weedy and generalist taxa (e.g., all three taxa of *Porites*, and massive taxa such as *Favia*, *Favites Platygyra, Lobophyllia*, and *Symphyllia*), which are tolerant and more adaptable to unfavorable conditions (Darling et al., 2012; Adjeroud et al., 2015; McWilliam et al. 2020). Overall, the study confirmed the advantages of inferring demographic processes from population size structure in hard corals. Using size-class frequency distribution analysis, in combination with common reef health indicators, the study provided a solid baseline for future monitoring of hard coral reef health at fine scale in Peninsular Malaysia, providing essential information for conservation assessments and management actions (Edmunds and Riegl, 2020). Ultimately, we suggest that hard coral demographic collapse is not a single-direction process. Much rather, structural demographic changes of taxa are subjected to a two-step filter. For instance, if taxa survive disturbances and are still fecund, whilst given a suitable degree of substrate availability, an initial shift towards recruitment and sub-adult dominated populations may be observed, accompanied by the elimination of non-tolerant and larger colonies susceptible to disturbances such as cyclones, overfishing and coral bleaching. If multiple disturbances persist in systems with reinforcing feedback loops (e.g., unconsolidated substrate coupled with monsoon waves), colony mortality and fragmentation rates will continue to increase and create unfavorable environments for demographic recovery and viability. We conclude that such reinforcing disturbance regimes are applying strong selective pressure on hard coral taxa, resulting in the survival of only the largest and most stress tolerant taxa with fewer functional traits, thus impoverishing the coral reef assemblage. Therefore, the reduction of the abundance of all size groups and a shift towards large colonies and weedy taxa may be the second step of a demographic continuum, where SFD of individual taxa is regionally determined by biophysical interactions (such as waves, colony shape and sedimentation), life-history traits, and the nature and persistence of stressor agents.

## Supporting information

Supplementary tables and figures (Bernard et al. 2022)

## Acknowledgments

We are thankful to Summer Bay Resort, Lang Tengah Island, for supporting our research with generous in-kind contributions. We are grateful to Albert Apollo Chan from the Department of Fisheries Malaysia (DoF) for endorsing the publication of this manuscript, under permit number Prk.ML.630-7Jld.5 (21). Special thanks is owed to Joseph A. Henry for providing critical suggestions that greatly improved the manuscript. The revision, writing and editing of this manuscript was supported with a research writing residency at Rimbun Dahan.

## Disclosure of funding

This research did not receive any specific grant from funding agencies in the public, commercial, or not-for-profit sectors. However, the researchers were supported in-kind by Summer Bay Resort, Lang Tengah Island, and received financial and non-financial support by Lang Tengah Turtle Watch and Coralku Solutions.

## Declaration of interests

The authors declare that they have no known competing financial interests or personal relationships that could have appeared to influence the work reported in this paper.

## Notes

### Competing Interest Statement

The authors have declared no competing interest.

