## Supplementary tables and figures (Bernard et al. 2022) for "Colony size frequency distribution across gradients of reef health in disturbed coral reefs in Northeast Peninsular Malaysia"

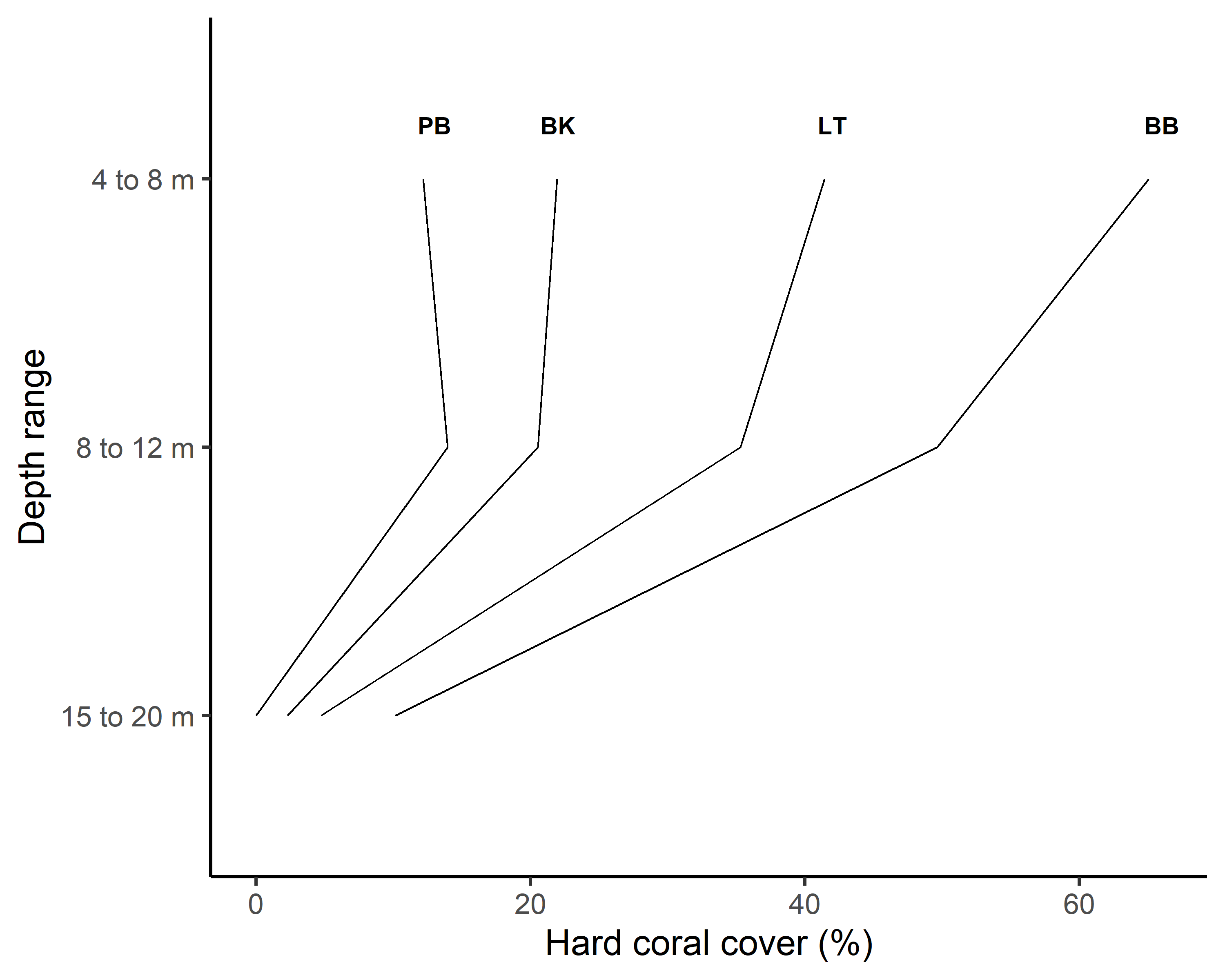

**Supplementary Figure S1 |** Percent live hard coral cover as a function of depth is presented for three sites at reef scale (BB-Batu Bulan; BK-Batu Kucing; PB-Pasir Besar) and for Pulau Lang Tengah (Pulah=Island) at 5°47’N, 102°53’E.

**Supplementary Table S2** | Summary of coral recruitment abundance across three sites at Pulau Lang Tengah (5°47’N, 102°53’E*)* in Northeast Peninsular Malaysia. Rec.% is the proportion of recruit colonies (< 5.0 cm) within each taxon, and Rel.% is each taxon’s relative recruitment abundance at reef-scale.

|  | Batu Bulan (BB) | | | | Batu Kucing | | | | Pasir Besar (PB) | | | |
| --- | --- | --- | --- | --- | --- | --- | --- | --- | --- | --- | --- | --- |
| Morpho-taxon | n | Density | Rec. % | Rel. % | n | Density | Rec. % | Rel. % | n | Density | Rec. % | Rel. % |
| All | 846 | 9.40 | 20.8 | 36.6 | 398 | 4.42 | 43.1 | 9.8 | 258 | 2.87 | 31.1 | 6.3 |
| *Acropora* corymbose | 10 | 0.11 | 40.0 | 0.5 | 15 | 0.17 | 26.7 | 1.0 | 126 | 1.40 | 1.6 | 0.8 |
| *Acropora* digitate | 11 | 0.12 | 0.0 | 0.0 | - | - | - | - | - | - | - | - |
| *Acropora* hispidose | 19 | 0.21 | 21.1 | 0.5 | - | - | - | - | 16 | 0.18 | 6.2 | 0.4 |
| *Acropora* hispidose-thick | - | - | - | - | - | - | - | - | 7 | 0.08 | 0.0 | 0.0 |
| *Cyphastrea* encrusting | 62 | 0.69 | 45.2 | 3.3 | 33 | 0.37 | 54.5 | 4.5 | - | - | - | - |
| *Diploastrea* massive | 9 | 0.10 | 22.2 | 0.2 | - | - | - | - | - | - | - | - |
| *Echinopora* encrusting | 54 | 0.60 | 22.2 | 1.4 | - | - | - | - | - | - | - | - |
| *Echinopora* laminar | 51 | 0.57 | 17.6 | 1.1 | - | - | - | - | - | - | - | - |
| *Favia* encrusting | 47 | 0.52 | 53.2 | 3.0 | 25 | 0.28 | 80.0 | 5.0 | - | - | - | - |
| *Favia* massive | 181 | 2.01 | 52.5 | 11.2 | 69 | 0.77 | 46.4 | 8.0 | 14 | 0.16 | 42.9 | 2.3 |
| *Favites* encrusting | 50 | 0.56 | 42.0 | 2.5 | 31 | 0.34 | 58.1 | 4.5 | - | - | - | - |
| *Favites* massive | 17 | 0.19 | 64.7 | 1.3 | 9 | 0.10 | 66.7 | 1.5 | - | - | - | - |
| *Fungia* solitary | 94 | 1.04 | 81.9 | 9.1 | - | - | - | - | 169 | 1.88 | 17.8 | 11.6 |
| *Galaxea* submassive | 207 | 2.30 | 20.8 | 5.1 | 56 | 0.62 | 41.1 | 5.8 | 7 | 0.08 | 57.1 | 1.6 |
| *Goniastrea* encrusting | 73 | 0.81 | 32.9 | 2.8 | 16 | 0.18 | 37.5 | 1.5 | - | - | - | - |
| *Goniastrea* encrustig-crumpled | 14 | 0.16 | 14.3 | 0.2 | - | - | - | - | - | - | - | - |
| *Goniastrea* massive | 14 | 0.16 | 28.6 | 0.5 | 13 | 0.14 | 46.2 | 1.5 | 7 | 0.08 | 100.0 | 2.7 |
| *Heliopora* columnar | 172 | 1.91 | 22.7 | 4.6 | - | - | - | - | - | - | - | - |
| *Leptastrea* encrusting | 70 | 0.78 | 68.6 | 5.7 | 33 | 0.37 | 75.8 | 6.3 | 22 | 0.24 | 77.3 | 6.6 |
| *Leptoria* massive | 9 | 0.10 | 0.0 | 0.0 | - | - | - | - | - | - | - | - |
| *Lobophyllia* massive | 28 | 0.31 | 57.1 | 1.9 | 7 | 0.08 | 28.6 | 0.5 | - | - | - | - |
| *Montipora* encrusting-upgrowth | 12 | 0.13 | 41.7 | 0.6 | - | - | - | - | - | - | - | - |
| *Montipora* encrusting | 11 | 0.12 | 36.4 | 0.5 | - | - | - | - | - | - | - | - |
| *Montipora* vase | 51 | 0.57 | 33.3 | 2.0 | 66 | 0.73 | 25.8 | 4.3 | - | - | - | - |
| *Pavona* encrusting | 15 | 0.17 | 66.7 | 1.2 | 14 | 0.16 | 71.4 | 2.5 | - | - | - | - |
| *Pavona* foliose | 36 | 0.40 | 33.3 | 1.4 | 10 | 0.11 | 30.0 | 0.8 | 66 | 0.73 | 33.3 | 8.5 |
| *Platygyra* encrusting | 32 | 0.36 | 40.6 | 1.5 | 10 | 0.11 | 70.0 | 1.8 | 7 | 0.08 | 42.9 | 1.2 |
| *Platygyra* massive | 114 | 1.27 | 21.1 | 2.8 | 33 | 0.37 | 18.2 | 1.5 | - | - | - | - |
| *Pocillopora* branching | 47 | 0.52 | 34.0 | 1.9 | 20 | 0.22 | 35.0 | 1.8 | 11 | 0.12 | 72.7 | 3.1 |
| *Porites* branching-thin | - | - | - | - | - | - | - | - | 23 | 0.26 | 65.2 | 5.8 |
| *Porites spp. (rus)* | 295 | 3.28 | 23.4 | 8.2 | 76 | 0.84 | 22.4 | 4.3 | 175 | 1.94 | 32.6 | 22.1 |
| *Porites* encrusting | 42 | 0.47 | 81.0 | 4.0 | 18 | 0.20 | 44.4 | 2.0 | - | - | - | - |
| *Porites* encrusting-crumpled | 11 | 0.12 | 18.2 | 0.2 | - | - | - | - | - | - | - | - |
| *Porites* massive | 288 | 3.20 | 35.8 | 12.2 | 291 | 3.23 | 42.6 | 31.2 | 95 | 1.06 | 46.3 | 17.1 |
| *Psammocora* branching | - | - | - | - | - | - | - | - | 17 | 0.19 | 29.4 | 1.9 |
| *Symphyllia* massive | 31 | 0.34 | 41.9 | 1.5 | 7 | 0.08 | 57.1 | 1.0 | - | - | - | - |
| *Turbinaria* encrusting | 8 | 0.09 | 62.5 | 0.6 | - | - | - | - | - | - | - | - |

| **Morpho-taxon** | **Site** | **n** | **Size_log_ (±SD)** | **10^th^** | **90^th^** | **g_1_ (Z)** | **g_2_ (Z)** | **CV** | |
| --- | --- | --- | --- | --- | --- | --- | --- | --- | --- |
| *Acropora* digitate | BB | 11 | 1.42 ± 0.18 | 1.18 | 1.48 | 0.02 (0.04) | -0.25 (-0.33) | | 12.76 |
| *Acropora* hispidose thick | PB | 7 | 1.69 ± 0.57 | 0.88 | 2.08 | -0.84 (-1.37) | -1.18 (-1.78) | | 33.62 |
| *Diploastrea* massive | BB | 9 | 1.29 ± 0.77 | 0.48 | 2.14 | 0.02 (0.03) | -1.2 (-1.63) | | 59.8 |
| *Echinopora* encrusting | BB | 54 | 1.24 ± 0.55 | 0.57 | 1.78 | -0.48 (-1.51) | -0.59 (-1.02) | | 44.56 |
| *Echinopora* laminar | BB | 51 | 1.19 ± 0.53 | 0.57 | 1.78 | 0.41 (1.26) | 0.37 (0.63) | | 44.23 |
| *Goniastrea* massive | BB | 14 | 1.13 ± 0.37 | 0.66 | 1.48 | 0.02 (0.04) | -1.06 (-1.36) | | 32.72 |
| *Heliopora* columnar | BB | 172 | 0.99 ± 0.40 | 0.57 | 1.48 | -0.07 (-0.36) | 0.07 (0.18) | | 40.47 |
| *Leptoria* massive | BB | 9 | 1.48 ± 0.21 | 1.18 | 1.78 | 0.00 (0.00) | -0.75 (-1.02) | | 14.41 |
| *Montipora* encrusting-upgrowth | BB | 12 | 0.95 ± 0.59 | 0.14 | 1.75 | 0.04 (0.07) | -1.22 (-1.58) | | 62.69 |
| *Montipora* encrusting | BB | 11 | 0.91 ± 0.47 | 0.57 | 1.48 | 0.20 (0.35) | -0.43 (-0.55) | | 51.56 |
| *Porites* thin branching | PB | 23 | 0.57 ± 0.33 | 0.10 | 0.88 | -0.26 (-0.57) | -0.96 (-1.3) | | 57.75 |
| *Porites* encrusting-crumpled | BB | 11 | 0.95 ± 0.47 | 0.10 | 1.48 | -0.92 (-1.61) | -0.28 (-0.36) | | 49.6 |
| *Psammocora* foliose | PB | 17 | 0.85 ± 0.38 | 0.38 | 1.18 | -0.63 (-1.25) | -0.15 (-0.19) | | 44.29 |
| *Turbinaria* encrusting | BB | 8 | 0.66 ± 0.56 | 0.10 | 1.27 | 0.26 (0.43) | **-1.48 (-2.10)** | | 84.48 |

**Supplementary Table S3 |** Colony size frequency distribution (SFD) statistics of morpho-taxa tested at one site (BB-Batu Bulan; BK-Batu Kucing; PB-Pasir Besar) around Pulau Lang Tengah in Northeast Peninsular Malaysia. Highlighted in grey are skewness and kurtosis of colony SFD distributions that are significantly different from the normal distribution.

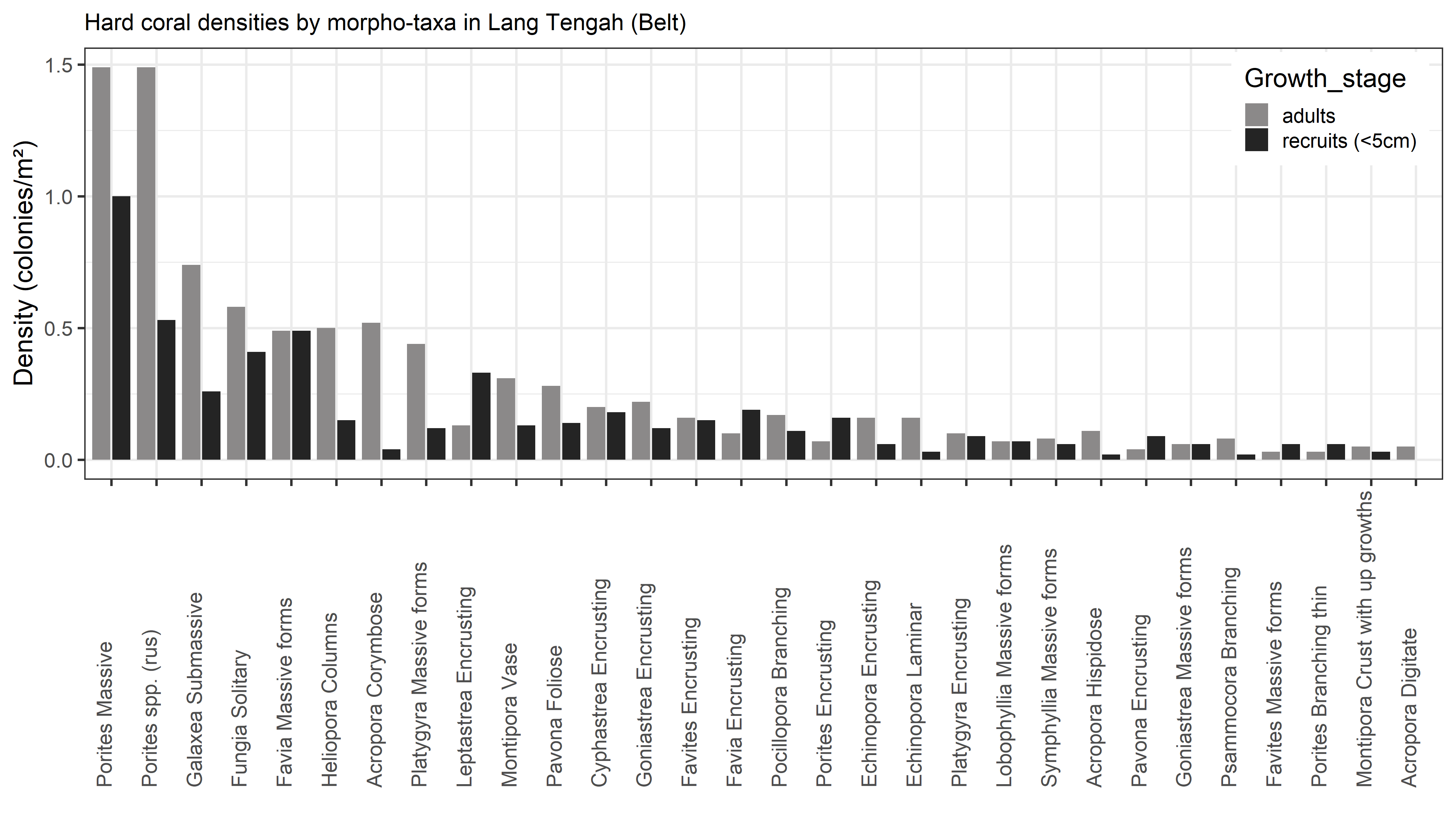

**Supplementary Figure S4 |** Rank abundance curve of hard coral morpho-taxa in Pulau Lang Tengah (5°47’N, 102°53’E) in Northeast Peninsular Malaysia as adult (black bars) and recruit (<5.0 cm, grey bars) colonies.

**Supplementary Table S5 |** List and percent relative abundance of hard coral morpho-taxa that cumulatively constitute two-thirds of the coral community assemblage on island (Lang Tengah) and reef scale, respectively.

| Morpho-taxon | Lang Tengah (LT) | Batu Bulan (BB) | Batu Kucing (BK) | Pasir Besar (PB) |
| --- | --- | --- | --- | --- |
| *Acropora c*orymbose | 3.50 | - | - | 15.20 |
| *Cyphastrea* encrusting | 2.44 | 2.89 | - | - |
| *Favia* massive | 6.78 | 8.45 | 7.50 | - |
| *Favites* encrusting | - | 2.33 | - | - |
| *Fungia* solitary | 6.76 | 4.39 | - | 20.39 |
| *Galaxea* submassive | 5.99 | 8.50 | 5.54 | - |
| *Goniastrea* encrusting | - | 3.41 | - | - |
| *Leptastrea* encrusting | 3.21 | 3.27 | 3.59 | - |
| *Montipora* vase | 3.01 | 2.38 | 7.17 | - |
| *Platygyra* massive | 3.78 | 5.32 | 3.59 | - |
| *Porites*massive | 17.32 | 13.45 | 31.63 | 11.46 |
| *Porites spp. (rus)* | 14.03 | 13.77 | 8.26 | 21.11 |
| **Total (%)** | **66.82** | **68.15** | **67.28** | **68.15** |

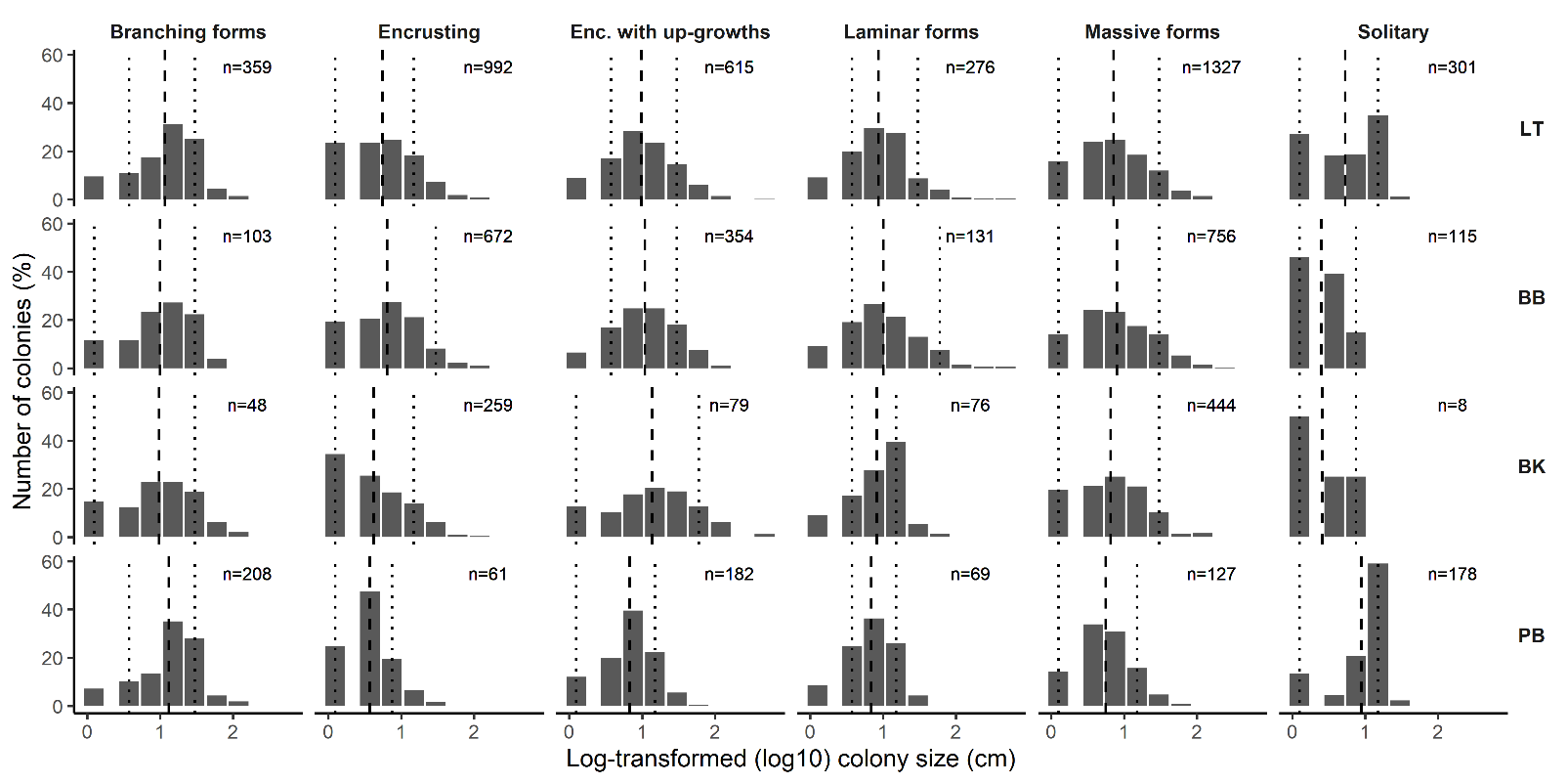

**Supplementary S6 |** Log-transformed size frequency distribution (SFD) of hard corals based on colony morphology at reef scale at three sites (BB-Batu Bulan; BK-Batu Kucing; PB-Pasir Besar) and on island scale (LT-Lang Tengah), in Northeast Peninsular Malaysia. Dotted lines present the 10^th^ and 90^th^ percentile, respectively, and the dashed line shows the mean of the distribution.
